# Spliceosome profiling visualizes the operations of a dynamic RNP *in vivo* at nucleotide resolution

**DOI:** 10.1101/224170

**Authors:** Jordan E. Burke, Adam D. Longhurst, Daria Merkurjev, Jade Sales-Lee, Beiduo Rao, James Moresco, John Yates, Jingyi Jessica Li, Hiten D. Madhani

## Abstract

Tools to understand how the spliceosome functions *in vivo* have lagged behind advances in its structural biology. We describe methods to globally profile spliceosome-bound precursor, intermediates and products at nucleotide resolution. We apply these tools to three divergent yeast species that span 600 million years of evolution. The sensitivity of the approach enables detection of novel cases of non-canonical catalysis including interrupted, recursive and nested splicing. Employing statistical modeling to understand the quantitative relationships between RNA features and the data, we uncover independent roles for intron size, position and number in substrate progression through the two catalytic stages. These include species-specific inputs suggestive of spliceosome-transcriptome coevolution. Further investigations reveal ATP-dependent discard of numerous endogenous substrates at both the precursor and lariat-intermediate stages and connect discard to intron retention, a form of splicing regulation. Spliceosome profiling is a quantitative, generalizable global technology to investigate an RNP central to eukaryotic gene expression.

**Highlights:**

- Measurement of spliceosome-bound precursor and intermediate in three species
- Non-canonical splicing events revealed
- Statistical modeling uncovers substrate features that predict catalytic efficiency
- Discard of suboptimal substrates occurs *in vivo* and predicts intron-retained mRNAs

## Introduction

A signature feature of eukaryotic gene expression is the splicing of primary transcripts by the spliceosome. Rather than serving as a mere impediment to gene expression, introns are central to gene regulation at many levels. In addition to controlling proteome diversity (Blencowe, 2017), splicing also impacts RNA stability, for example through the inclusion of exons with premature termination codons subject to nonsense-mediated decay (NMD) (Amrani et al., 2006; Brogna and Wen, 2009; Luo et al., 2001; Maquat, 2004; Wiegand et al., 2003; Zhou et al., 2000). Splicing also plays critical roles in RNA export and translation efficiency (Luo et al., 2001; Maquat, 2004; Wiegand et al., 2003; Zhou et al., 2000). Biogenesis of noncoding RNA often requires splicing, including many snoRNAs and miRNAs that are encoded within introns rather than exons (Hesselberth, 2013). Finally, splicing is a major player in human disease; a large fraction of single nucleotide polymorphisms associated with human disease impact splicing (Li et al., 2016), and many human cancers harbor driver mutations in components of the spliceosome itself (Bejar, 2016).

Our understanding of the assembly dynamics of the spliceosome comes almost entirely from decades of studies of model intron substrates in splicing-competent cell extracts from *S. cerevisiae* and HeLa cells (Wahl et al., 2009), and has been substantially further illuminated by recent high-resolution structural work (Fica and Nagai, 2017; Shi, 2017). Activation of the spliceosome after its assembly involves major ATP-dependent rearrangements that expel two RNA-protein complexes, the U1 and U4 snRNPs, and several proteins (Wahl et al., 2009). At the same time, the Prp19 complex (NTC) and NTC related proteins (NTR) join the complex to produce the B^act^ complex. These events trigger assembly of the RNA catalytic core of the spliceosome. Splicing then proceeds via two transesterification reactions. In step 1, nucleophilic attack of the 2’ hydroxyl of an adenosine at the branchpoint (BP) on the phosphate at the 5’ splice site (ss) produces a cleaved 5’ exon and branched species harboring a 2’-5’ phosphodiester bond called the lariat intermediate (Figure 1A). The second transesterification is achieved through attack of the 3’ hydroxyl of the cleaved 5’ exon on the phosphate at the 3’ss, producing the ligated exons and excising the intron as a branched RNA, referred to as the lariat intron. A handful of additional factors are required to bind and then leave the spliceosome during these two steps. These include the DEXD-box ATPases Prp16 and Prp22, which are thought to remodel/inactivate the active site after the first and second chemical steps, respectively, enabling intermediates and products to proceed to the next step in mRNA biogenesis (Krishnan et al., 2013; Ohrt et al., 2013; Schwer, 2008; Semlow et al., 2016; Tseng et al., 2011). Following mRNA release, the excised lariat intron and associated snRNPs are dissociated by the helicase Prp43 (Martin et al., 2002; Tsai et al., 2005). The 2’-5’ phosphodiester bond at the BP of the lariat intron is hydrolyzed by the debranching enzyme, Dbr1 (Chapman and Boeke, 1991; Ruskin and Green, 1985).

**Figure 1.**
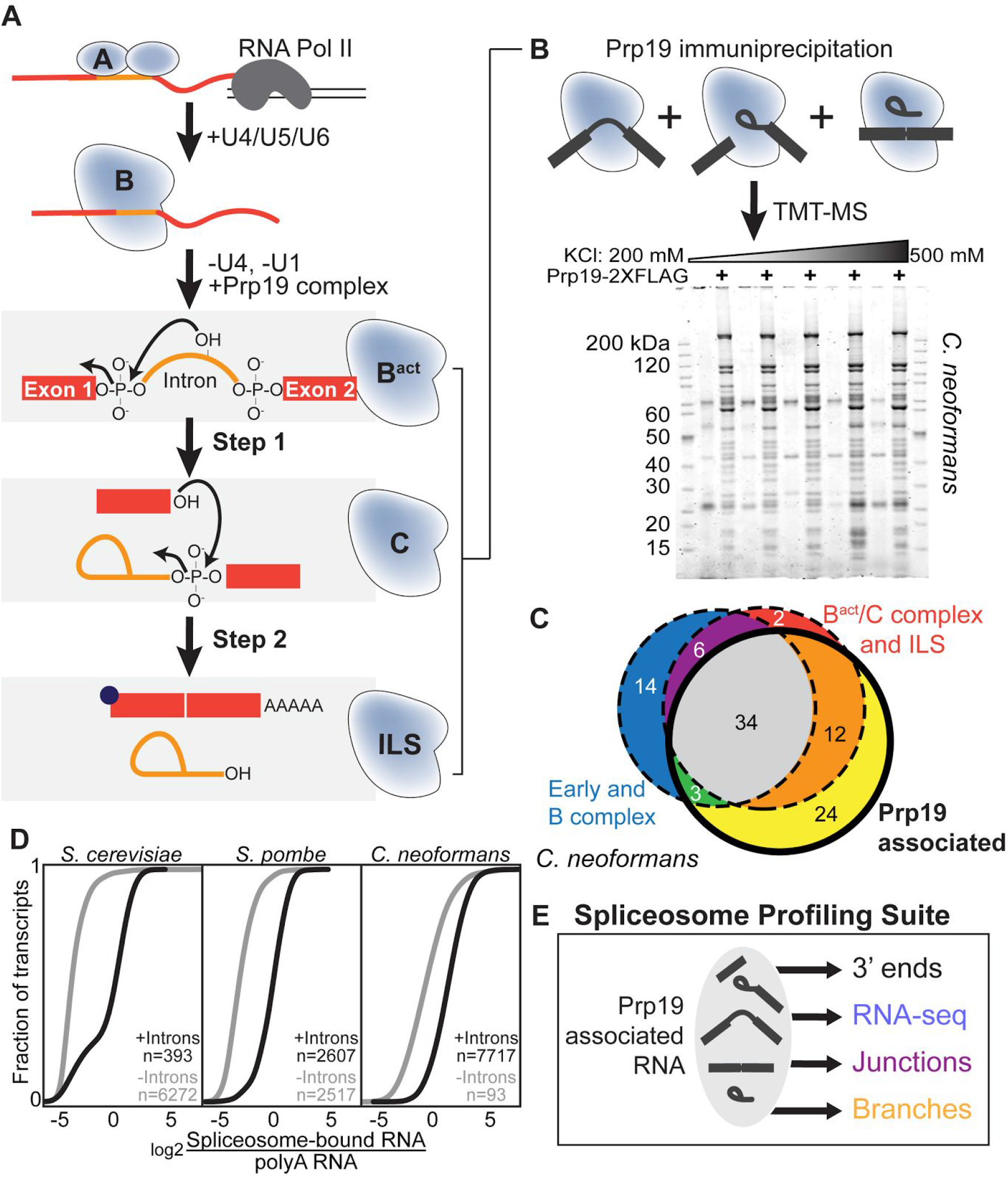
Isolating RNA from purified active spliceosomes using Prp19. A. Simplified view of the spliceosome assembly pathway and catalytic cycle indicating relevant substrate transformations and ribonucleoprotein complexes. B. SDS-PAGE (4–12% Acrylamide Tris-Glycine, Novex) analysis of spliceosomes affinity-purified from *C. neoformans* extracts using FLAG-tagged Prp19 at increasing KCl concentrations. Proteins were visualized using SYPRO Ruby. C. TMT-MS reveals that Prp19 associated splicing factors belong primarily to B^act^, C and ILS complexes. Membership of *S. cerevisiae* orthologs in complexes were used to assign factors. The 24 factors in the yellow portion of the Venn diagram are orthologs of splicing factors found in *H. sapiens* and/or *S. pombe* but not *S. cerevisiae* (See Figure S1B for list) as well as 6 peptidyl prolyl isomerases. Likely contaminants (ribosomal proteins and metabolic enzymes) are not shown. D. RNAs associated with Prp19 spliceosomes are enriched for transcripts containing annotated introns (KS test, p-values: *S. cerevisiae*: 3X10^-154^, *S. pombe*: < 2X10^-308^, *C. neoformans*: 9X10^-16^). E. RNA from Prp19-associated spliceosomes is used for multiple purposes in the spliceosome profiling suite.

The spliceosome is thus both remarkably dynamic and complex. These properties are thought to promote the fidelity of splicing through kinetic proofreading schemes, while permitting flexibility and regulation. Studies using *S. cerevisiae* splicing-competent extracts have shown that mutant pre-mRNA substrates that assemble into spliceosomes but are kinetically slow at either chemical step trigger spliceosome disassembly prior to completion of the reaction, a process termed “discard.” Together with prior suppressor genetics and associated biochemical work, this work has led to a model for spliceosomal fidelity in which the failure to perform catalysis prior to ATP hydrolysis by Prp16 (step 1) or Prp22 (step 2) produces a spliceosome “marked” for discard, which signals disassembly by Prp43 (Koodathingal et al., 2010; Mayas et al., 2010; Mayas et al., 2006; Semlow et al., 2016).

While these studies have been important for our understanding of the mechanism of fidelity, they are based on studies of mutant RNAs in a single species, *S. cerevisiae*, that harbors few introns and these display unusually strong matches to 5’ss, BP and 3’ss consensus signals (Irimia and Roy, 2008; Kupfer et al., 2004). The match of each signal to consensus positively impacts progression of an intron through the steps of splicing (Lesser and Guthrie, 1993). Fungi from other subphyla, such as *S. pombe* and *C. neoformans*, are considerably more intron-rich and display a much broader range of splicing signals, akin to those of plants and animals (Duff et al., 2015; Irimia and Roy, 2008; Kupfer et al., 2004; Sibley et al., 2015; Suzuki et al., 2013). Because of these considerations, it remains unclear to what extent inefficiency triggers discard in native introns *in vivo* and whether discard contributes significantly to the regulation of the RNA population as a whole, particularly in intron populations with diverse sequences at the 5’ss and BP regions.

RNA-seq, the high-throughput sequencing of cDNAs from fragmented mRNA, has enabled genome-wide identification of splicing events *in vivo*. Although the sensitivity of RNA-seq is limited by RNA stability, high-depth experiments can identify rare sequencing reads corresponding to splicing precursors and products. Even non-canonical splicing events such as recursive splicing involving exons of size zero have been identified in mammals and Drosophila (Duff et al., 2015; Sibley et al., 2015; Suzuki et al., 2013). Mutation of factors involved in nuclear or cytoplasmic RNA turnover or surveillance mechanisms can enable detection of splicing events that produce highly unstable RNAs, but these mutations are confounding as they necessarily alter the RNA population. Likewise, excised lariats can be stabilized in cells containing a knockout of *DBR1* (Awan et al., 2013; Gould et al., 2016; Mayerle et al., 2017), but this approach has the same drawback. Sequencing of chromatin-associated RNA has been used to enrich for shorter-lived products of splicing (Bhatt et al., 2012; Mayer et al., 2015), but this method requires such products to co-fractionate with the genome. Lariat-introns have been enriched by immunoprecipitation of *S. cerevisiae* extracts with polyclonal antibodies against Prp16 (Qin et al., 2016); this qualitative method enriches for a subpopulation of spliceosomes transiently bound by this ATPase. Despite these advances, none of these approaches directly interrogates the full population of transcripts bound to active spliceosomes, nor can they be used to quantify the progression of substrates through the spliceosome cycle.

We describe here a strategy for purification of the full complement of endogenous active spliceosomes using a cross-organism tagging and purification strategy followed by high throughput sequencing library preparation schemes to identify total spliceosome bound RNA, step 1 and step 2 cleavage sites as well as methods to quantify relative precursor and intermediate levels. We apply it to three highly divergent yeasts, the brewing yeasts *S. cerevisiae* and *S. pombe* and the human pathogenic yeast *C. neoformans*, which diverged from a common ancestor approximately 600 million years ago (Lucking et al., 2009; Prieto and Wedin, 2013). Below we describe these methods, which we term the spliceosome profiling suite, their utility for investigating the spliceosome *in vivo*, and the insights gained.

## Results

### Prp19 is a species-independent handle for isolating active spliceosomes

The spliceosome is not a single entity, but rather a series of dynamic complexes (Figure 1A). Based on *in vitro* assembly studies, the NTC component Prp19 joins the spliceosome during the process of activation and is associated primarily with *S. cerevisiae* spliceosomes that are activated and bound to mRNA precursor (B^act^), intermediate (C), or product (ILS) (Figure 1A) (Fabrizio et al., 2009). Affinity purification of Prp19-TAP from *S. pombe* also yields components diagnostic of these complexes (Ren et al., 2011). To extend these studies to *C. neoformans*, we affinity-purified Prp19-FLAG associated spliceosomes from cell extracts prepared at a range of salt (KCl) concentrations (Figure S1A) and performed tandem mass tag mass spectrometry (TMT-MS) to quantify the purified proteins (Figure 1B). We find that while orthologs of most annotated splicing factors copurify with Prp19 in lysates prepared in buffers containing up to 500 mM KCl (Figure 1B and Table S1), 55 splicing components exhibit quantitatively decreased abundance (based on reporter ion intensities) at KCl concentrations at and above 300 mM. Because copurification of common contaminants is also decreased at KCl concentrations above 300 mM KCl (Figure 1B and Table S1), we chose the 300 mM KCl condition for *C. neoformans*. Based on titration experiments, we found that endogenous spliceosomes in *S. cerevisiae* and *S. pombe* are somewhat less salt-stable; thus, 200 mM KCl was used for these yeasts (data not shown).

Of splicing factor orthologs that co-purify with Prp19 in *C. neoformans*, 49/72 are orthologs of proteins that are part of the B^act^, C and ILS complexes of *S. cerevisiae* (Figure 1C and Table S1), indicating that the Prp19 purification effectively enriches for actively splicing and post-catalytic spliceosomes. An additional 24 factors associated with *C. neoformans* Prp19 have no orthologs in *S. cerevisiae* and all but 2 have human orthologs annotated as spliceosome-associated factors (Cvitkovic and Jurica, 2013). These include GPATCH1, HNRNPUL1, DHX35 and members of the exon junction complex (Figure S1B and Table S1), which are associated with human B^act^ and C complex (Bessonov et al., 2010).

### Profiling spliceosome-associated RNA

We extracted RNA from affinity-purified spliceosomes from *S. cerevisiae, S. pombe* and *C. neoformans* using endogenously tagged Prp19 alleles (TAP in *S. cerevisiae* and *S. pombe* and FLAG in *C. neoformans*) and a single-step purification protocol (Figure S1A). We also isolated total RNA from whole-cell extracts and selected polyadenylated (polyA) RNA to measure the abundance of spliceosome-bound RNA and mature RNA. Comparing the abundance of the transcript associated with the spliceosome to the abundance of the polyA mRNA, we observed that, in all three yeasts, Prp19-associated RNA is enriched for transcripts having at least one annotated intron (Figure 1D). As detailed below, RNA associated with Prp19-spliceosomes was processed, sequenced and analyzed in a number of different ways to illuminate the action of the spliceosome *in vivo* (Figure 1E).

### 3’ end profiling of transcripts reveals intermediates and products of splicing

Genome-wide methods to analyze the rate or efficiency of splicing, such as splicing microarrays (Clark et al., 2002) and high-throughput sequencing (Brooks et al., 2011; Katz et al., 2010; Shen et al., 2014) typically rely on measuring precursor and spliced RNA. In contrast, using RNA from spliceosomes, we could directly observe the spliceosome-bound precursors, intermediates and products of the splicing reaction globally. To detect the intermediates and products, we labeled free 3’ hydroxyl groups using an approach adapted from NET-seq (Figure 2A) (Churchman and Weissman, 2011). In this protocol, a 5’-adenylated DNA linker is ligated to available RNA 3’ OH groups, allowing cDNA to be subsequently synthesized from the 3’ end of the RNA via priming from the DNA linker. The cDNA is then circularized, PCR-amplified, and subject to sequencing. Importantly, we include a 6 nt random barcode at the end of the DNA linker, allowing the computational detection and collapse of sequencing reads corresponding to a single ligation event to prevent over-counting of PCR amplified DNA species. In addition to profiling 3’ ends, we also prepared spliceosome-bound RNA and polyA RNA from whole cell extract for standard RNA-seq analysis (Figure 2A). For these samples, we first hydrolyzed the RNA by heating under alkaline conditions and then repaired the 3’ phosphate with T4 polynucleotide kinase as previously described for ribosome profiling (Ingolia et al., 2012). We then reverse-transcribed the RNA, circularized and amplified the resulting cDNA to produce sequencing libraries using the same steps as 3’ end profiling.

**Figure 2.**
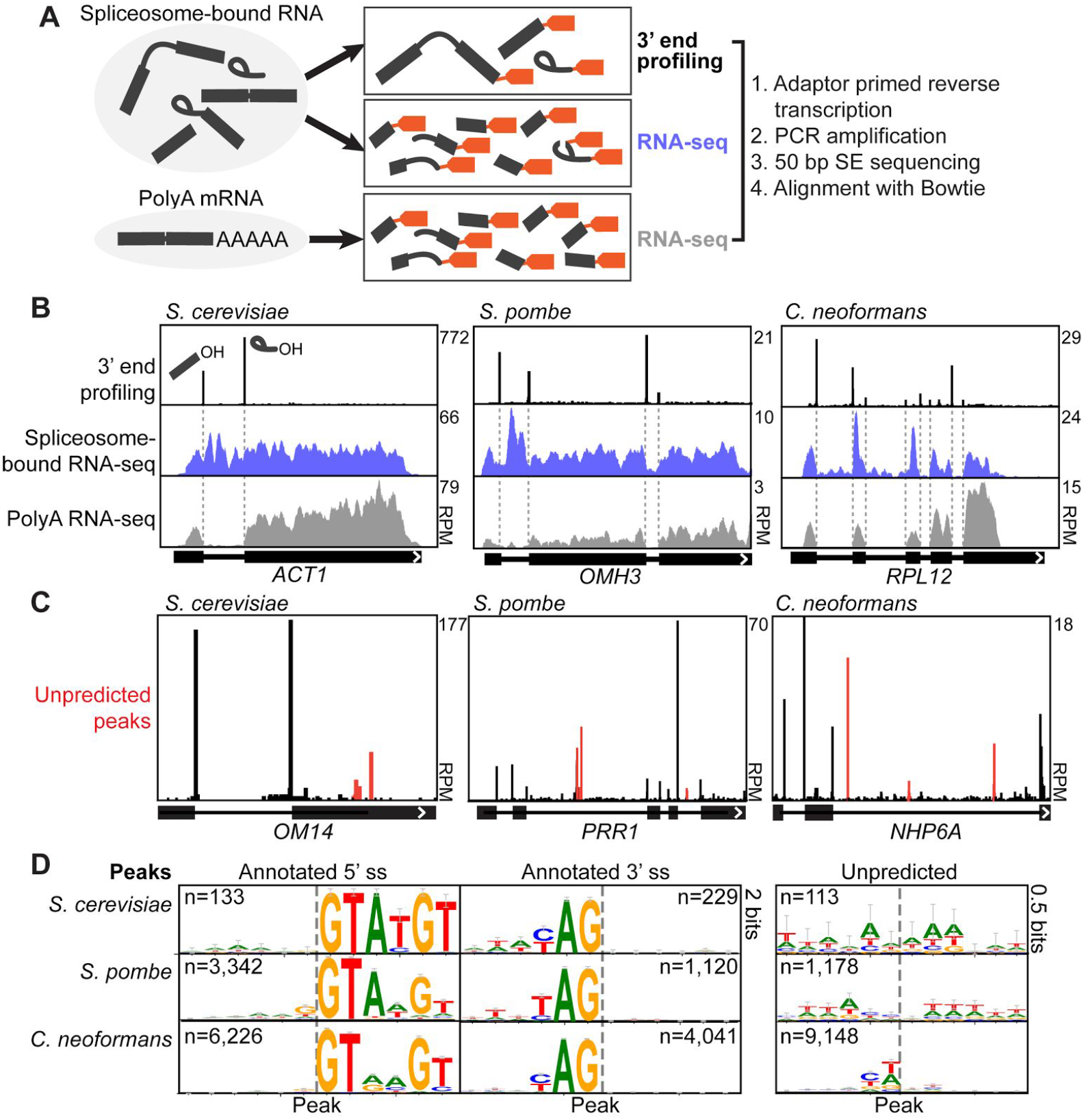
3’ end profiling reveals splicing intermediates and products in three yeasts. A. Strategies for 3’ end profiling and RNA-seq of spliceosome-bound and polyA RNA (see Results and Methods for details). B. 3’ end profiling (black traces) reveals splicing intermediates (cleaved 5’ exon) and products (lariat intron) globally in the three indicated organisms. Spliceosome-bound (blue) and polyA (grey) RNA-seq coverage for each transcript are also shown. C. 3’ end profiling also discovers novel 3’ ends (red) that may be spliceosomal cleavages or transcript ends. D. Logos from predicted and unpredicted peaks detected by 3’ end profiling in each of the three species.

In each of the yeasts, we observed pile-ups of reads beginning at annotated 5’ splice sites that correspond to 3’OH of the cleaved 5’ exon, an intermediate of the splicing reaction (Figure 2B). We also observe pile-ups of reads at annotated 3’ss whose ends correspond the 3’ OH of the excised lariat intron, a product of the reaction (Figure 2B). Comparing the RNA-seq data from spliceosome-bound RNA to polyA mRNA, we observe substantially more reads in introns, consistent with enrichment for unspliced transcripts and excised intron. 3’ end intensities at annotated splice sites and read density across spliceosome-bound and polyA RNA are reproducible between biological replicates (Figure S2A-F shows data for the intron-rich species, *S. pombe* and *C. neoformans*).

To automatically detect peaks, we developed a pipeline using the PELT (Pruned Exact Linear Time) method (Killick et al., 2012) in the R package changepoint (Killick and Eckley, 2014). We applied this approach (see Methods) to two replicate experimental datasets obtained using a tagged Prp19 allele and data from an untagged control experiment. Peaks that were detected in both experimental datasets but not in the control were retained for analysis. We observed peaks corresponding to annotated splice sites, as well as peaks throughout transcripts (Figure 2C). While the sequences neighboring peaks at annotated sites mirror the consensus sequence for each organism as expected, peaks corresponding to unannotated sites did not follow splicing consensus sequences (Figure 2D, right panel). The majority of these may correspond to non-adenylated/uridylated transcript 3’ ends produced by pausing or termination of RNA polymerase II or endonucleolytic cleavage. Only a minority of reads contributes to peaks in this category (6.5% in *S. cerevisiae*, 7.2% in *S. pombe* and 22% in *C. neoformans*).

### Junction and branch profiling confirm identities of 3’ end peaks

To identify the subset of unpredicted peaks that correspond to *bona fide* cleaved 5’ exons at unannotated 5’ splice sites and or excised intron lariats at unannotated 3’ splice sites, we took advantage of existing tools for detecting the complementary species produced by the splicing reaction, namely the branch formed after the first step of splicing or the exon-exon junction formed after the second step (Figure 3A). To observe branching events, we enriched for lariat-intermediate and lariat-intron species by treating spliceosome-bound RNA with the 3’-5’ exonuclease, RNase R (Suzuki et al., 2006). We then synthesized cDNA under conditions that enable read-through of lariats (see Methods), constructed sequencing libraries and performed 100 bp paired-end sequencing. After aligning reads to the genome with Tophat (Trapnell et al., 2009), we searched those that did not align to the genome for reads that begin downstream of a 5’ss or unpredicted peak and end downstream inside the same intron (Figure 3A). To identify exon-exon junctions, we prepared cDNA suitable for sequencing using random priming and performed 100 bp paired-end sequencing on spliceosome-bound RNA, using TopHat to detect exon-exon junctions. Of junctions detected in RNA-seq analysis of spliceosome-bound RNA in *C. neoformans*, 34% were observed only in spliceosome-bound RNA and not in polyA RNA (Figure 3B). In contrast, only 1% of junctions observed by analysis of polyA RNA were not observed in spliceosome-bound RNA (Figure 3B). Thus, analysis of junctions in spliceosome-associated RNA enables the detection of thousands of events that do not lead to the accumulation of polyadenylated RNA, presumably because such products are subject to decay mechanisms.

**Figure 3.**
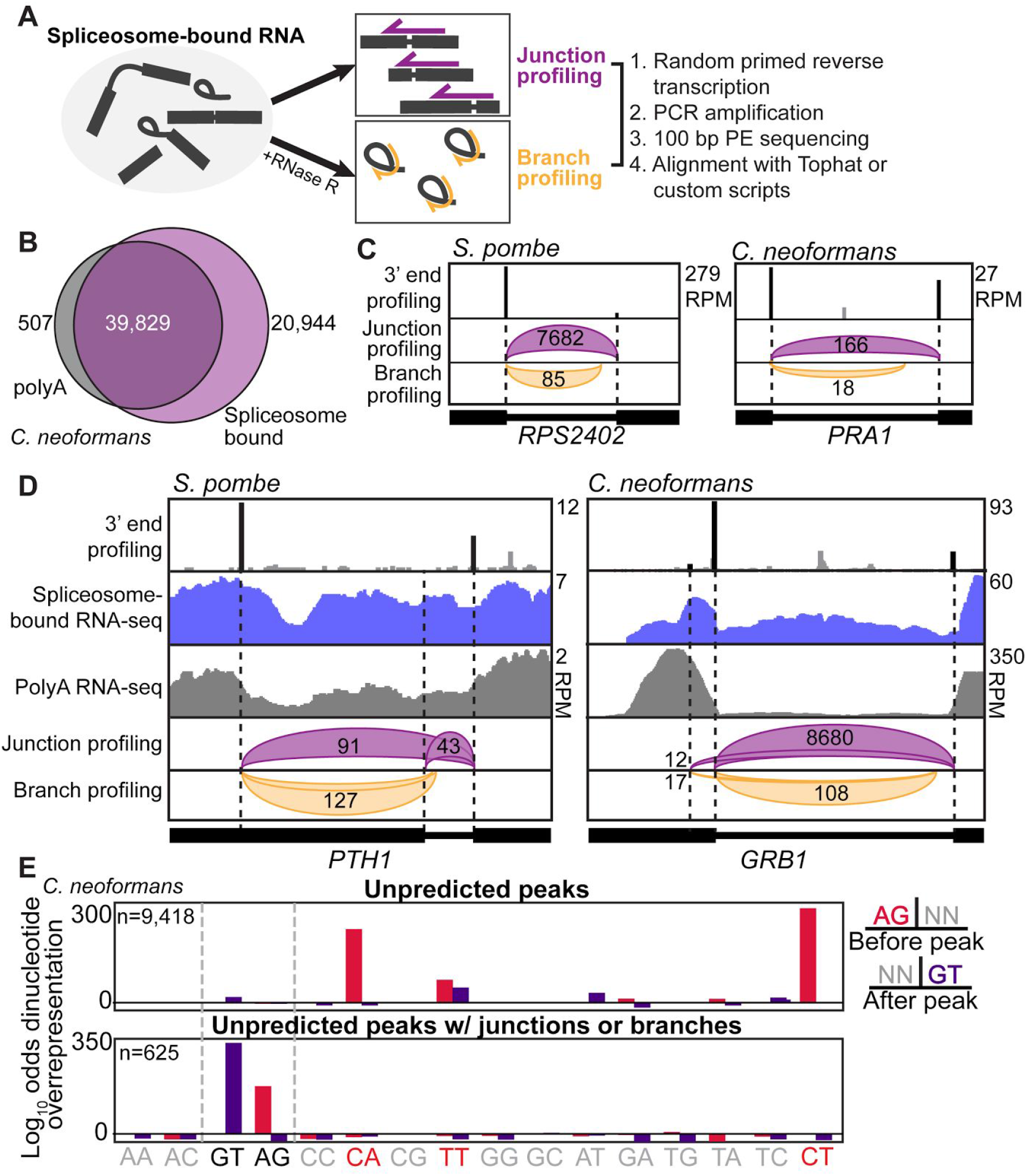
Branch and junction profiling confirm splicing events discovered by 3’ end profiling. A. Schematic of junction and branch profiling library preparation. Magenta arrows indicate reads that span exon-exon junctions and curved yellow arrows indicate reads that span branches. B. Overlap of junctions discovered by polyA vs. spliceosome-bound RNA-seq using Tophat. C. Example of an exon-exon junction and branch detected at an annotated introns in *S. pombe* and *C. neoformans* (*PRA1*: CNAG_04950). Purple and yellow curves are derived from junction BED files generated by Tophat (junction profiling) or custom scripts (branch profiling) visualized in IGV. Numbers indicate number of reads covering each exon-exon junction or branch (not normalized). D. Examples of junctions and branches that confirmed unannotated alternative 5’ splice sites in *S. pombe* and *C. neoformans* (*GRB1*: CNAG_03281). Peaks that align with other splicing events are shown in black. Other 3’ end profiling signals are shown in grey. E. Dinucleotide overrepresentation (as determined by log(odds ratio)) immediately upstream and downstream of peaks in *C. neoformans* (see Figure S3B for *S. pombe)*. Unfiltered unpredicted peaks are enriched for pyrimidine rich dinucleotides and splicing signal dinucleotides while peaks with a junction or branch are only enriched for splicing signal dinucleotides.

To select high-confidence splicing events, we developed a pipeline that compares branches and junctions to peaks detected by 3’ end profiling (see Figure 3C for examples, algorithm design in Figure S3A). Using this approach, we identified thousands of peaks that correspond to annotated and unannotated complementary splicing events in *S. pombe* and *C. neoformans*. Figure 3D shows examples of alternative 5’ss choice in *S. pombe* and *C. neoformans*. We observe 89 unannotated 5’ or 3’ splice sites in *S. pombe* and 446 in *C. neoformans* that have a detectable peak by 3’ end profiling and either a junction or branch. Within these high-confidence sets, dinucleotides immediately before the peaks are strongly enriched for the GT dinucleotide (5’ss) and the dinucleotides immediately after the peaks are strongly enriched for the AG dinucleotide (3’ss), consistent with the identification of *bona fide* splicing events in both *C. neoformans* (Figure 3E) and *S. pombe* (Figure S3B).

### Spliceosome profiling detects non-canonical events

We next tested whether the sensitivity of spliceosome profiling enabled the detection of unusual splicing events. Using junction profiling, we detected numerous alternative splicing events (738 in *S. pombe* and 1367 in *C. neoformans*) that coincide with at least one peak from 3’ end profiling. Within these unannotated events, we searched for examples of types of non-canonical splicing. One such type is interrupted splicing, where the spliceosome recognizes a 5’ss and BP and performs the first step but then does not proceed to the second step due to lack of an acceptable splice receptor (Figure 4A). In *S. cerevisiae*, interrupted splicing of the *BDF2* transcript (encoding a BET family bromodomain protein) results in degradation of the transcript by the nuclear exosome in a process called spliceosome mediated decay (Volanakis et al., 2013). Indeed, we observed a substantial amount of spliceosome-bound intermediate for the *BDF2* transcript and no peaks that correspond to a 3’ss (Figure 4B). In *S. pombe*, the 3’ end of the telomerase RNA, *TER1*, is formed by the first step of splicing occurring without the second (Kannan et al., 2013). We observed the intermediate formed by this reaction (Figure 4C), but also the exon-exon junction to a nearby 3’ss; however, transcripts with this junction appear to be unstable as only the full length *TER1* transcript was detected in polyA mRNA (Figure 4C, grey trace) and the relative read coverage of the junction is low compared to that of canonically spliced transcripts (Figure S4A). We also uncovered one other transcript subject to interrupted splicing, an unannotated *S. pombe* transcript named SPBC530.07c (Figure 4D and S4A). This gene encodes a member TENA/THI-4 family of proteins, which includes enzymes involved in thiamine metabolism (Akiyama and Nakashima, 1996; Pang et al., 1991). The *BDF2, TER1* and SPBC530.07c transcripts are enriched 30-75 fold on the spliceosome relative to unspliced transcripts (Figure S4B-C), consistent with accumulation of intermediates on the spliceosome.

**Figure 4.**
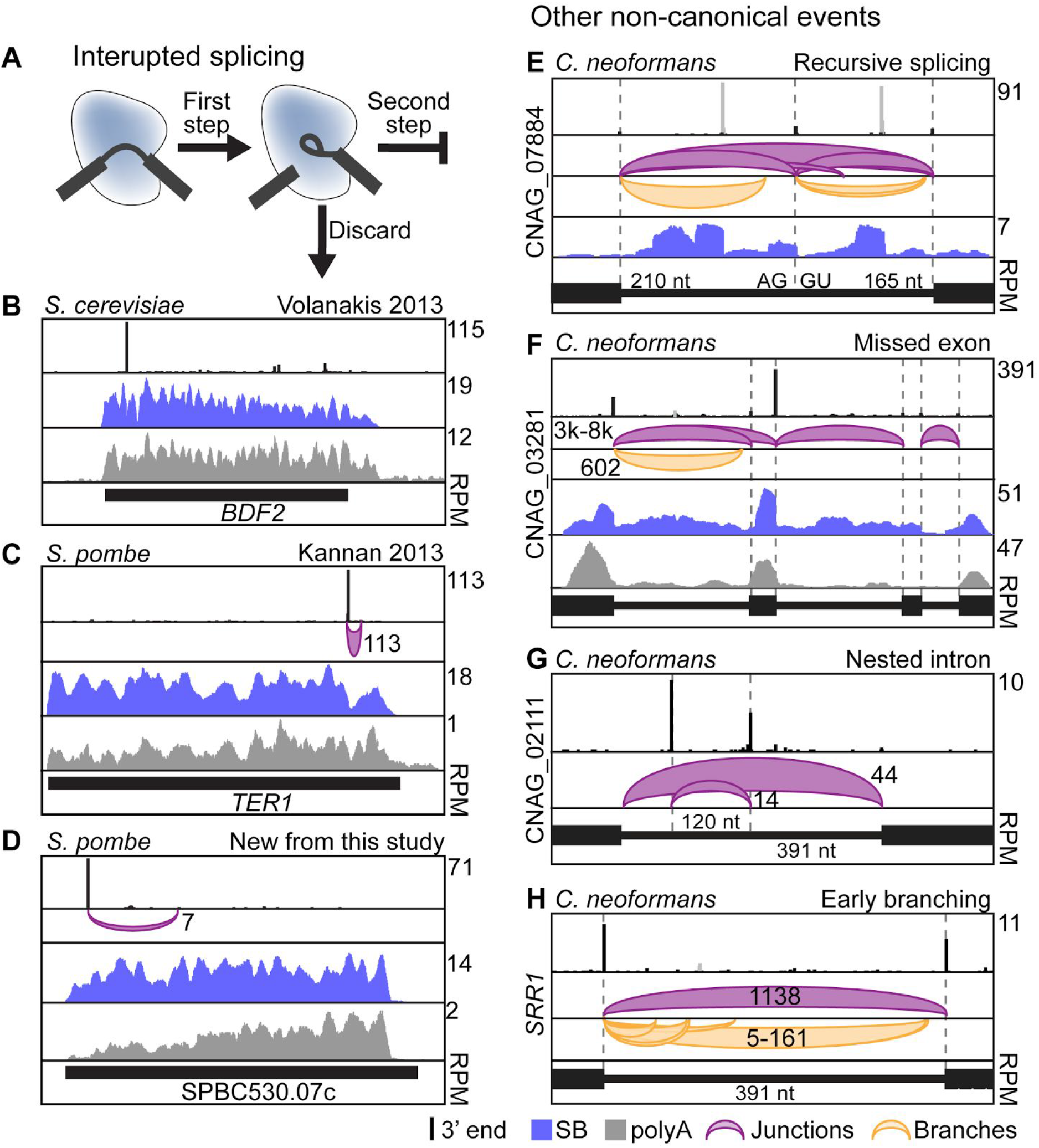
Non-canonical splicing events. A. Splicing can be interrupted between the two steps when discard is triggered by a slow second step (Kannan et al., 2013; Volanakis et al., 2013). B-D. Examples of interrupted splicing. 3’ end profiling results are shown in black, spliceosome-bound RNA-seq data are shown in blue, polyA RNA-seq data are shown in grey and junction profiling results are shown in purple. The number of reads corresponding to each junction (not normalized) is indicated. E. An example of recursive splicing in *C. neoformans*, where an annotated intron is actually two introns separated by a zero nucleotide exon “pivot point.” Traces are colored as above and branch profiling is shown in yellow. Grey peaks correspond to putative 3’ RNA ends based on the profile of the spliceosome-bound RNA-seq data for this transcript. F. An example of a splicing event resulting in exclusion of an exon in *C. neoformans*. Traces colored as above. G. Nested intron inside a relatively large (391 nt) intron. Only the 391 nt intron is detectable by polyA RNA-seq (data not shown). Traces colored as above. H. An example of early branches inside a relatively large (391 nt) intron. In these cases, the branch to 3’ss distances are much larger than the typical range of 15-30 nt. Traces colored as above.

In addition to interrupted splicing, we observed other non-canonical phenomena. This includes recursive splicing, the removal of introns that flank a zero-nucleotide exon, which has recently been reported to be extensive in *D. melanogaster* (Duff et al., 2015) and in mammals (Sibley et al., 2015) based on intron coverage patterns and junctions in high-depth RNA-seq data. We identified a recursive splicing event in *C. neoformans* in a 375 nt intron of the *BOT1/CNAG_07884* gene, which encodes an protein essential for *C. neoformans* viability (Ianiri and Idnurm, 2015) related to the *S. pombe* Bot1 mitochondrial ribosomal protein (Wiley et al., 2008). Multiple lines of evidence indicate that recursive splicing effectively splits this intron into two smaller ones of 210 and 165 nt (Figure 4E). Specifically, we identified a peak that includes signal from both the lariat intron of the first splicing event and the free 5’ exon of the second splicing event. We also identified junctions and branches for both events (Figure 4E, magenta and yellow, shorter arches) and the junction that corresponds to the final exon-exon junction (Figure 4E, magenta, long arch). We observed two additional large peaks (Figure 4E, grey) that do not correspond to splicing events but appear to be the 3’ ends of premature transcripts based on spliceosome-bound RNA-seq (Figure 4E, blue trace). In another transcript, an alternative 3’ss is chosen that, curiously, results in the precise skipping of the downstream exon (Figure 4F). The isoform lacking this exon was not detected in the mature polyA mRNA (data not shown).

Another type of non-canonical splicing is nested splicing where a smaller intron is excised from a larger intron potentially reducing the size of the intron (“intron within an intron”). We observed 13 examples of nested splicing in *C. neoformans* (one example is shown in Figure 4G). We also identified many instances (63 in *S. pombe* and 104 in *C. neoformans)* of introns where branch sites unusually distal from the 3’ss are recognized and used for the first step (Figure 4H). These “early” or “distal” branching events have also been detected in RNA-seq data from human cells (Taggart et al., 2017). In both *S. pombe* and *C. neoformans*, they are particularly common in introns larger than 150 nt (Figure S4D). In *C. neoformans*, 43% of early branching events result in BP-3’ss distances greater than 50 nt (Figure S4E), which may be prohibitively large for the second step of splicing.

### Estimating the efficiency of splicing with spliceosome profiling

Assuming steady state kinetics, reproducible measurements of the relative abundance of pre-mRNA and intermediate bound to Prp19-associated spliceosomes (normalized to total spliceosome-bound RNAs) would enable estimates of the ability of endogenous substrates to progress through the first and second steps *in vivo*, thereby providing insights into how substrates behave once activated spliceosomes have assembled.

To accomplish this, we focused on high-confidence splicing events identified by our pipeline by first defining a transcript set that is above a threshold for the quantification limit of the assay (at least one high-confidence splicing event within each transcript; *S. cerevisiae*: 128, *S. pombe*: 1889, *C. neoformans*: 2121). Within this set, we quantified spliceosome-bound precursor and intermediate levels at each annotated intron and any unpredicted 5’ splice sites. We defined the level of bound precursor as the number of unique reads that begin at least 5 nt downstream of the 5’ss and end at least 5 nt upstream (Figure 5A) normalized to the total read density of spliceosome-bound RNA across the transcript (Figure 5B). Based on the assumption that the steady-state level of precursor bound to the spliceosome would be inversely proportional to the rate of the first step of splicing, we used precursor level to estimate the relative efficiency of the first step of splicing. We defined the level of bound intermediate as the number of unique reads that begin at the 5’ss (corresponding to the cleaved 5’ exon, Figure 5A) normalized to the total read density of spliceosome-bound RNA across the transcript (Figure 5B) and we employed this ratio to estimate the relative efficiency of the second step of splicing. Spliceosome-bound precursor and intermediate level measurements were both reproducible between biological replicates, with intermediate level measurements displaying higher reproducibility (Figure 5B and S5A-B). Precursor levels form a narrower distribution in all three organisms than intermediate levels (Figure 5B and S5A-B, Levene test (log2 transform) p-values – *S. cerevisiae*: 2 X10^-22^, *S. pombe*: 3 X10^-279^, *C. neoformans*: 5 X10^-295^).

**Figure 5.**
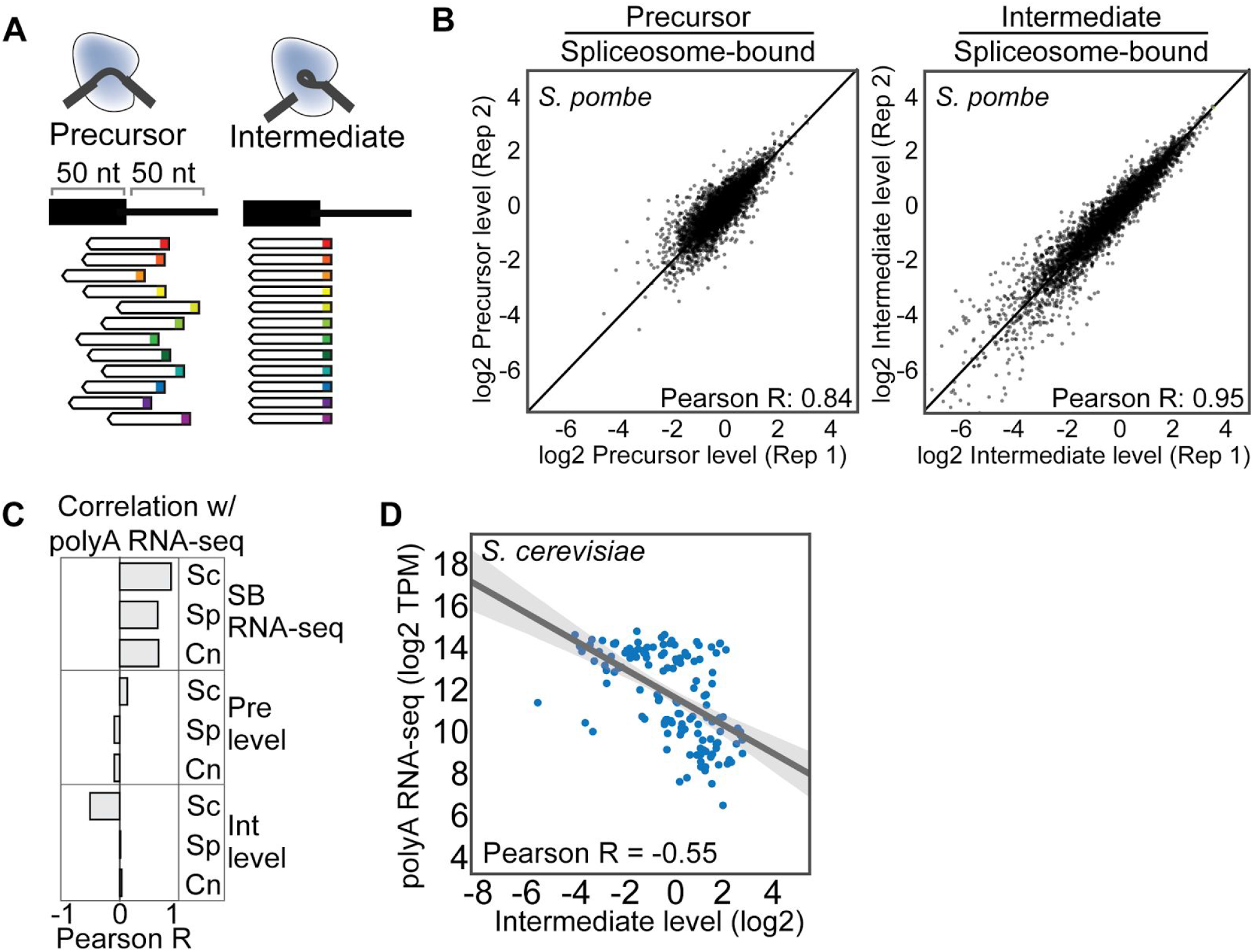
Quantitation of spliceosome-bound precursor and intermediate. A. Quantitating the level of precursor and intermediate at each splice site. Precursor is determined by counting reads that start downstream of the splice site and end upstream. Intermediate is determined by counting reads that begin at the splice site. Each read begins with a random hexamer bar code (rainbow colors). B. Reproducibility and distribution of precursor and intermediate levels (precursor and intermediate normalized to the read density of spliceosome-bound transcript) in *S. pombe* (see Figure S5A-B for other yeast). C. Correlation of precursor, intermediate and spliceosome-bound transcript levels with the abundance of the mature transcript as determined by polyA RNA-seq (log2 transformed to approximate normality for Pearson R analysis). D. Correlation of polyA RNA-seq with intermediate levels in *S. cerevisiae*. Grey area indicates 95% confidence interval for the linear regression model.

The total amount of transcript associated with Prp19 spliceosomes positively correlates with the abundance of polyA mRNA in the cell as determined by RNA-seq (Figure 5C) and is likely related to the rate of transcription. In contrast, mRNA abundance does not positively correlate with spliceosome-bound precursor or intermediate level in any yeast, suggesting a strong role for substrate differences in controlling progression through the spliceosome cycle *in vivo*. Unexpectedly, relative intermediate levels on spliceosomes from *S. cerevisiae* are strongly negatively correlated with mRNA abundances (Figures 5C and D).

### Bayesian modeling identifies features that predict bound precursor and intermediate levels

Because spliceosome profiling provides the opportunity to assay splicing efficiency across thousands of substrates, we sought to exploit the statistical power of the sample sizes to investigate, in an unbiased fashion, intron features that predict the levels of normalized spliceosome-bound precursor and intermediate. As any such approach is data-driven, we restricted our analysis to the intron-rich species *S. pombe* and *C. neoformans*. We employed Bayesian Model Averaging, a probabilistic approach that generates large numbers of linear models, averages the best models and determines the probability, magnitude and uncertainty of the contribution of a given feature to the final ensemble of models (Clyde, 2017). The probability of including an intron feature in the final model is expressed as the marginal posterior inclusion probability or PIP (see Methods). A PIP of 0.5 is interpreted as an equal probability (50%) that a feature is significantly predictive of the response variable. In our case, the response variable is the level of normalized spliceosome-bound precursor or intermediate. We tested a range of features based on our current understanding of introns and the spliceosome (Figure S6A and Table S6). We focus here on three aspects of the results of this analysis: 1) the correlation of match to intron consensus with spliceosome-bound precursor and intermediate, 2) the correlation of distance-related features such as intron size, and 3) roles of intron number and position.

We observed PIPs of 1.0 for a negative correlation of 5’ss and BP scores with spliceosome-bound precursor levels in both *S. pombe* and *C. neoformans*, presumably reflecting the role of these sequences in step 1 catalysis (Figure 6A-D and S6A-B), such that a stronger 5’ss and BP results in more rapid transition through the first step (or the immediately preceding steps). 5’ss, BP, pyrimidine tract, and 3’ss scores are positively correlated with intermediate levels in *S. pombe* and BP and pyrimidine tract scores in *C. neoformans* (PIPs=1.0, Figure 6B-C and S6B-C). The predictive roles of these sequences for intermediate levels may reflect higher accumulation of the products of step 1 catalysis (the cleaved 5’ exon and lariat-intermediate) when splicing signals are more optimal. They could also reflect reduced rates of spliceosomal remodeling steps required to execute the second chemical step (see Discussion).

**Figure 6.**
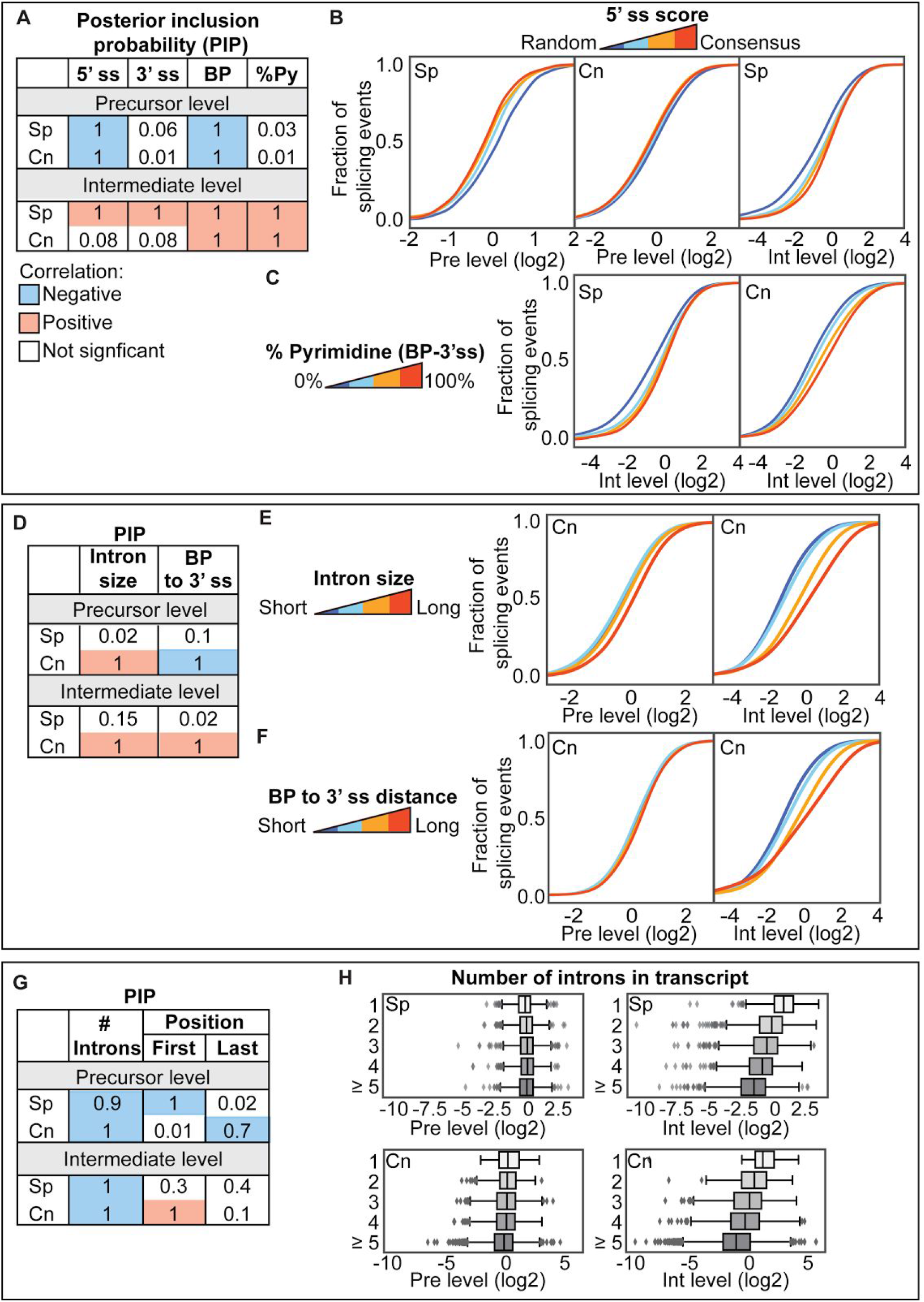
The relationship between intron features and splicing efficiency. A. Posterior inclusion probabilities (PIPs) for Bayesian Model Averaging of the relationship between intron features and precursor and intermediate levels. PIPs for 5’ss, 3’ss and BP scores and % pyrimidine between the BP and 3’ss are shown. Intron features with a PIP higher than 0.5 are highlighted (red = positive correlation, blue = negative correlation). See Figure S6A for coefficients and uncertainties. B. Relationship between precursor and intermediate levels and 5’ss scores. Introns were split into quartiles based on 5’ss score (blue=random sequence, red=consensus) and the distribution of each metric is plotted as a CDF (See Figure S6B-C for BP and 3’ss scores). Sp indicates *S. pombe* and Cn indicates *C. neoformans*. C. Same as B but for the percent pyrimidine in the region between the branch point and the 3’ ss. D. PIPs for intron size and the distance between the BP and 3’ss. Intron features with a PIP higher than 0.5 are highlighted (red = positive correlation, blue = negative correlation). E. Relationship between precursor and intermediate levels and intron length in *C. neoformans*. Introns were split into quartiles based on length (blue=short, red=long) and the distribution of each metric for each quartile is plotted as a CDF. F. Same as E but for BP to 3’ss distance. G. PIPs for the number of introns in the transcript and the relative position of the intron in the transcript. Intron features with a PIP higher than 0.5 are highlighted (red = positive correlation, blue = negative correlation). H. Difference in precursor and intermediate levels based on the number of introns in the transcript (See Figure S6F-G for alternative splicing and position in transcript).

Strikingly, a predictive feature of both precursor and intermediate levels (PIPs=1.0) in *C. neoformans* identified by the models is intron size (Figure 6D-E and S6A). This feature is not predictive in *S. pombe* (PIP=0.02 and 0.15, for precursor and intermediate). Notably, *C. neoformans* introns display a size distribution that is significantly narrower than that of *S. pombe* (Figure S6D). Precursor and intermediate levels are contributed to by the distance between the BP to 3’ss in *S. pombe* and *C. neoformans* (PIPs=1.0; Figure 6F), and are also positively predicted by the presence of detectable alternative BP or 3’ss utilization events (PIP=1.0, Figure S6A and F). We also identified sizes of upstream and downstream exons as species-specific predictors (Figure S6A). Taken together, these data indicate that the distance relationships between substrate signals play a role in the efficiency of splicing after spliceosome assembly *in vivo* in a species that displays a relatively narrow intron length distribution.

Finally, we found that the number of introns in the transcript and the position of the intron in the transcript are also predictive of precursor and intermediate levels (Figure 6G-H). Transcripts with more introns predict lower intermediate levels in both *S. pombe* and *C. neoformans* (PIPs=1.0, Figure 6G and H, bottom panels). Precursor levels are also negatively predicted by intron number (PIPs=0.9-1.0, Figure 6G-H). Introns that are first in the transcript also exhibit higher intermediate level in *C. neoformans*, but lower precursor and intermediate level in *S. pombe* (Figure 6G and S6A and G).

### Intron retention may be triggered by discard after spliceosome assembly

Intron retention is a major form of alternative splicing that is generally thought to be the result of inefficient spliceosome assembly. An untested alternative possibility for the source of such intron-retained species derives from the kinetic proofreading model of splicing fidelity described in the Introduction, in which suboptimal pre-mRNA substrates can be discarded from the spliceosome *after* assembly due to slow catalysis (Koodathingal and Staley, 2013). This hypothesis predicts a correlation between the levels of intron retention in the polyA RNA population and the normalized levels of spliceosome-bound precursor, as high levels of the latter reflect slower progression through first-step catalysis after assembly.

We measured intron retention in RNA-seq data produced from polyA RNA, defining the level of retention as the read density inside the intron normalized to the total read density across the transcript. In *S. cerevisiae*, we observed no significant correlation between spliceosome-bound precursor level and intron retention (Figure 7A, left panel). However in the other two yeasts, which display denser and more diverse intron populations, we observed significant correlations, indicating that precursor molecules that progress slowly through the first chemical step after spliceosome assembly are more likely to be retained in the polyA population (Figure 7A). These data are consistent with the model that *S. pombe* and *C. neoformans* execute discard after assembly to produce intron-retained species in the polyA population.

**Figure 7.**
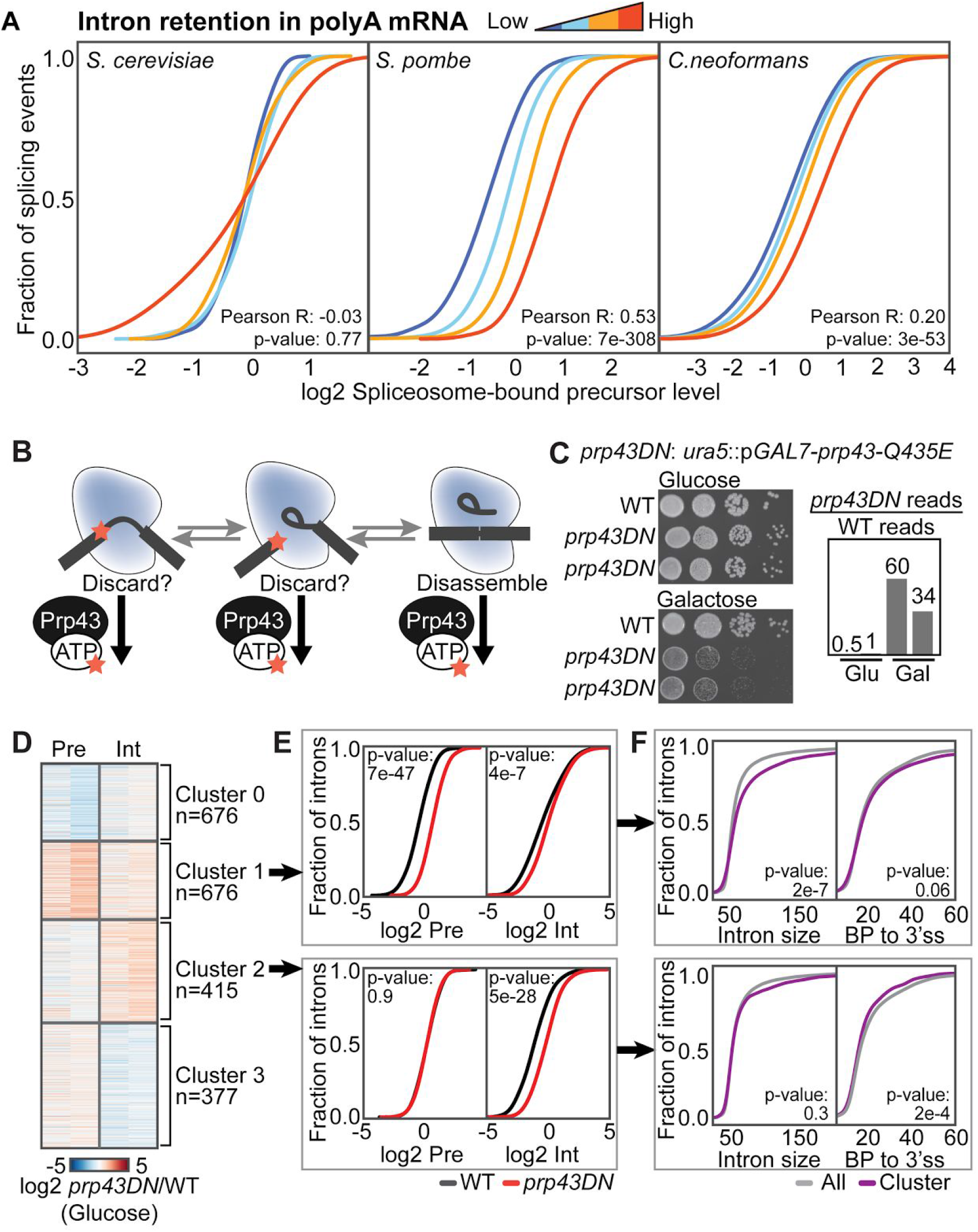
Prp43 discards long introns before the first step of splicing. A. Intron retention correlates with precursor level in *S. pombe* and *C. neoformans* but not *S. cerevisiae*. Introns were split into quartiles based on the level of intron retention calculated in polyA RNA-seq data. The distribution of precursor level in each quartile is plotted as a CDF (See Figure S7A for intermediate level). B. Simplified model of Prp43 mediated discard from the spliceosome (adapted from (Koodathingal and Staley, 2013)). This model predicts that disabling the ATPase activity of Prp43 will cause precursor and intermediates to accumulate on spliceosomes. C. Growth of strains expressing *prp43-Q435E* or *prp43DN* from the *URA5* locus in glucose or galactose media and relative expression of *prp43-DN* (compared to the wild type transcript) in each condition, determined by determining the number of reads with and without the corresponding nucleotide change from polyA RNA-seq data. D. Heat map of the ratio of precursor and intermediate in the *prp43DN* strains vs. wild type grown under repressive conditions. Introns are sorted by K-means clustering with 4 centroids (see Methods). E. CDF plots of precursor and intermediate in *prp43DN* (red) and wild type (black) in glucose for cluster 1 (top panel) and cluster 2 (bottom panel) where intermediate or precursor, respectively, is increased in the *prp43DN* strains. P-values determined by KS test. F. Intron size and BP to 3’ss distance for each cluster compared to all introns for cluster 1 (top panel) and cluster 2 (bottom panel). P-values determined by Mann-Whitney U test.

Unexpectedly, we also observed similar correlations with intermediate level (Figure S7A); however, in this case, *S. cerevisiae* and *C. neoformans* exhibited the strongest effects. This correlation may occur because defects in signals that result in slow step 1 kinetics also result in slow step 2 kinetics – this is known to be the case for certain splice site mutations in *S. cerevisiae* (Lesser and Guthrie, 1993; Liu et al., 2007; Query and Konarska, 2004).

### Prp43 mediated discard of precursor and intermediate species *in vivo*

Nonetheless, the result that inefficiently spliced introns are more likely to appear as retained introns in the polyadenylated mRNA population generally supports the hypothesis that substrates can be discarded from the spliceosome after its assembly (Mayas et al., 2010). To further test this idea, we employed a mutation in the RNA helicase that disassembles the spliceosome upon successful completion of splicing, Prp43 (Tsai et al., 2005). *In vitro* studies in *S. cerevisiae* have demonstrated that addition of a dominant-negative form of Prp43 that binds the spliceosome but cannot hydrolyze ATP results in accumulation of spliceosome-bound lariat intermediate that would otherwise be released from spliceosomes via its disassembly (Mayas et al., 2010). These studies utilized a substrate mutated at the 3’ss to block the second catalytic step. We hypothesized that expression of this allele would also lead to accumulation of precursor and intermediate species of endogenous introns on spliceosomes *in vivo* if they were normally subjected to discard and Prp43-dependent disassembly (Figure 7B). To test this hypothesis, we created a strain of *C. neoformans* that expresses the equivalent dominant negative Prp43 allele, *prp43-Q435E* (hereafter referred to as *prp43DN)* under control of the galactose-inducible *GAL7* promoter inserted at the *URA5* locus. Under repressive glucose media conditions, the *prp43DN* transcript accumulates to roughly the same abundance as the wild type transcript (Figure 7C), indicating leaky expression of the *GAL7* promoter, but the strain grows as well as wild type on glucose (Figure 7C). Under inducing galactose media conditions, the *prp43DN* transcript accumulates to 30-60 times the level of the wild type transcript and the strain displays a marked growth defect (Figure 7C). No splicing factors exhibited decreased expression in either condition (see RNA-seq data in Figure S7B and Table S7); nonetheless, to minimize potential indirect effects, we performed spliceosome profiling analysis on wild type and the *prp43DN* strain cultivated in glucose-containing media. As with the analyses described above, experiments were performed in duplicate.

To examine changes in the steady-state spliceosome-bound substrate populations in the *prp43DN* strain, we grouped introns based on normalized bound precursor and intermediate levels using K-means clustering (Figure 7D, see Methods). Of these clusters, one exhibits a significant increase in spliceosome-bound precursor levels (Cluster 1, Figure 7D-E) and another exhibits a significant increase in spliceosome-bound intermediate levels in the *prp43DN* strain (Cluster 2, Figure 7D-E). These data support the model that Prp43 disassembles spliceosomes *in vivo* prior to completion of the catalytic steps and that specific sets of introns are subject to this activity.

Introns whose spliceosome-bound precursor and intermediate levels are controlled by Prp43 do not display significantly weaker 5’ss or 3’ss sequences (Figure S7C-D). Rather, introns with increased intermediate levels in the *prp43DN* strain display shorter BP to 3’ss distances on average (Figure 7F) and detectably stronger 5’ss and BP scores (Figure S7E-F, BP scores – Mann-Whitney U p-value: 4 X10^-6)^. Introns in this cluster are also more likely to be the last intron (Figure S7E). In contrast, introns with increased spliceosome-bound precursor in the *prp43DN* background are significantly longer (Figure 7F). Introns in this cluster are also more likely to be first introns (Figure S7E). Together with the studies of intron retention above, these studies provide evidence for widespread discard of natural spliceosome-bound substrates and reveal endogenous predictors of this behavior.

## Discussion

The spliceosomal layer of gene expression determines the amount and structure of proteins and ncRNAs produced in the cell. Understanding these mechanisms during normal homeostasis and disease requires tools to interrogate this RNA-protein machine *in vivo*. We have described a suite of spliceosome profiling tools that marry the biochemical purification of endogenous spliceosomes with high-throughput sequencing/analysis methods to identify high-confidence splicing events in spliceosome-bound RNA, quantifying both the precursor and intermediates as well as identifying products that evade detection by RNA-seq. By implementing these methods in three widely divergent yeast species, we demonstrate that the method can be established in multiple systems. Although we used tagged alleles of the essential splicing factor Prp19, which is located at the periphery of all reported high resolution spliceosome structures, in principle, any accessible tag or epitope could be used, making the approach adaptable to any organism for which sufficient numbers of cells can be obtained. Spliceosome profiling has substantial advantages over current methodologies because 1) it quantifies spliceosome-bound precursor and intermediate levels, 2) it identifies trans-esterification events that cannot be easily identified by RNA-seq because of their transient nature or decay after release from the spliceosome, 3) it does not require additional mutations to stabilize RNA species. Consequently, we anticipate its widespread application for the analysis of conditions, drugs, and mutations that impact gene expression via the spliceosome. Below we discuss insights gained from this initial study and prospects for further development.

### Integration of data reveals novel spliceosome-catalyzed transesterification events

Our initial analysis of RNA 3’ ends associated with spliceosome-bound transcripts revealed cleavages at the boundaries of annotated introns as well as numerous unpredicted ends. Because many of these ends do not appear to be produced by the action of the spliceosome, we filtered these data using information gained from profiling spliceosome-bound branchpoints and junctions. Through this approach, we obtained a high-confidence set of previously unannotated spliceosome-catalyzed events in *S. pombe* and *C. neoformans*. We detected non-canonical events including interrupted splicing events in *S. cerevisiae* and *S. pombe* including those involved in spliceosome-mediated decay and the biogenesis of telomerase RNA. We also identified a number of nested splicing events (“introns within introns”) in *C. neoformans*. Finally, we identified an essential *C. neoformans* gene, *BOT1*, that is subject to recursive splicing in *C. neoformans*. Such events have previously only been described in animal cells for very long introns (Duff et al., 2015; Sibley et al., 2015). We anticipate that at least some of these non-canonical events will be verified as targets for regulation.

### Quantification enables investigation of substrate features that influence progression

We proceeded beyond qualitative identification of cleavage and ligation sites by focusing on spliceosome-bound substrates sufficiently abundant to be quantified reproducibly. This enabled searches for the determinants of substrate behavior in fully assembled spliceosomes *in vivo*, which was not possible with prior technologies. The much larger number of quantifiable splicing events in the intron-rich yeast species *S. pombe* and *C. neoformans* allowed for the use of an automated statistical modeling approach to identify predictive substrate features. A clear finding was that stronger 5’ss and BP sequences predict lower bound precursor levels in *S. pombe* and *C. neoformans*, as expected for a role for the intron itself in catalytic efficiency (or immediately preceding conformational changes). This finding is consistent with recent cryo-EM structures that show known and novel intimate interactions between the 5’ss and BP sequences with the catalytic core of the spliceosome (Fica and Nagai, 2017; Shi, 2017). Surprisingly, more optimal intron sequences (5’ss, BP, pyrimidine tract and 3’ss) predicted higher intermediate levels, particularly in *S. pombe*. As mentioned above, this may reflect the more rapid accumulation of intermediate as a result of the effects of more optimal sequences on the first chemical step.

Modeling also revealed predictive roles for splice site and BP spacing and spliceosome-bound precursor and intermediate levels in *C. neoformans*, but not *S. pombe*. As mentioned above, this is fully consistent with the tighter distribution of intron sizes in *C. neoformans*, which has been shown to be under complex evolutionary selective forces (Hughes et al., 2008). Consistent with prior work, we observe that long introns in *S. pombe* and *C. neoformans* tend to harbor detectably stronger splicing signals (Figure S6C), further suggesting evolutionary pressure to compensate for non-optimal intron size. Our observation of ‘early branching’ in long introns may be indicative of size being measured prior to the first chemical step. The corresponding longer distance between the BP and 3’ss could then impact the second chemical step as this is a well established determinant of step 2 kinetics in *S. cerevisiae* (Luukkonen and Séraphin, 1997; Zhang and Schwer, 1997).

Numerous species display narrow intron length distributions, with the most remarkable being the ciliate *Stentor* in which nearly all introns are precisely 15 nucleotides in length (Slabodnick et al., 2017). The *Drosophila* genome is enriched for introns of 60-65 nucleotides in length (Lim and Burge, 2001) and smaller introns may be selected for during evolution due to improved splicing efficiency (Carvalho and Clark, 1999). Intron length has been shown to impact overall efficiency during *in vitro* splicing in HeLa and in *Drosophila* cell extracts (Fox-Walsh et al., 2005; Guo and Mount, 1995) and *in vivo* (Ulfendahl et al., 1985) but whether this is due to an impact on spliceosome assembly or catalysis has not been assessed; our studies demonstrate a role after assembly *in vivo*. Identifying the yardstick that measures intron size in assembled spliceosomes in species that display such constraints will likely require structural analysis.

Our modeling studies also revealed that higher numbers of introns in a transcript correlate with lower levels of bound intermediate in both *S. pombe* and *C. neoformans*. An intriguing explanation for this correlation would be that spliceosomes bound to nearby introns on a transcript can influence each other’s activities. Testing these and other hypotheses inspired by the modeling will require further investigation.

### Spliceosome discard of native transcripts

The ability of spliceosomes stalled *in vitro* to be disassembled is well established. However, as these studies require alterations to the substrate that induce stalling, the extent to which such events occur frequently in normal cells and what their function might be is unclear. Spliceosomal discard has been conceptualized as a mechanism for enhancing the fidelity of splicing by limiting the impact of splicing errors. This thinking is based on substantial *in vivo* and *in vitro* analysis of mutant substrates in *S. cerevisiae*. While a reasonable inference, it has not been tested in the context of naturally-occurring transcripts, in part because deviation from consensus is rare in *S. cerevisiae*, but also because tests for discard *in vivo* have been challenging to develop. A notable exception is the finding that Prp43 is important for the biogenesis of the *TER1* RNA in *S. pombe* via interrupted splicing (Kannan et al., 2013). For this transcript, discard appears to promote a specific biogenesis pathway, rather than a mechanism for the suppression of errors. Our finding that intron retention correlates with high levels of spliceosome-bound precursor suggests another role for discard in the biogenesis of intron-retained RNA, a common form of alternative splicing. Intron retention has recently been shown to be regulated by environmental stresses in *C. neoformans* (Gonzalez-Hilarion et al., 2016) raising the possibility that there exist corresponding signal transduction mechanisms that impinge on fully assembled spliceosomes.

In complementary studies, we identified precursor and intermediate populations that accumulate on spliceosomes in cells expressing ATPase-defective version of Prp43. This finding provides the *in vivo* evidence for Prp43 activity upstream of spliceosome disassembly on a substantial fraction of native transcripts prior to the completion of splicing. Our analysis revealed several predictors of this behavior (intron size for Prp43-dependent precursor discard and BP to 3’ss distance for Prp43-dependent intermediate discard), suggesting that these introns are more likely to undergo discard under normal conditions. Notably, these substrates did not differ in the overall strength of splicing signals. Additional parameters may determine the sensitivity of these transcripts to Prp43 activity.

### Prospects

The suite of spliceosome profiling tools described here was implemented in three divergent yeasts, demonstrating the adaptability of the methods. The ability to interrogate mutants in spliceosomal components combined with the data analysis approaches described here offers an attractive approach for understanding the function of spliceosomal components/protein domains/residues, including many whose three-dimensional location is now known but whose precise function across substrates remains to be dissected. Our proteomic characterization of spliceosomes in *C. neoformans*, a species that harbors over 40,000 annotated introns, indicates it harbors numerous factors that have human orthologs but are not found in the intron-reduced species *S. cerevisiae*. These include components of the exon junction complex. Analysis of these factors by spliceosome profiling may illuminate mechanisms important for species harboring a higher degree of splicing complexity. Finally, translation of our protocols to mammalian cells should enhance studies of the role of the spliceosome in human disease.

## Supplemental material

### Supplemental tables

**Table S1.** Mass spectrometry results with values, related to Figure 1

**Table S2.** Prp19 transcript enrichment for spliced and unspliced transcripts, related to Figure 1

**Table S3.** All peaks detected for each organism, related to Figure 2

**Table S4.** All detected branches and junctions for *S. pombe* and *C. neoformans*, related to Figure 3

**Table S5.** Cross-confirmed peaks for each organism with quantitation, related to Figure 3

**Table S6.** Results of Bayesian model averaging, related to Figure 6

**Table S7.** Differential expression analysis of polyA RNA-seq for *prp43DN* samples, related to Figure 7

**Table S8.** Filtered precursor and intermediate level measurements for *prp43DN*, related to Figure 7

**Table S9.** Summary of experiments and deposited processed and raw data files.

### Supplemental figure legends

**Figure S1.**
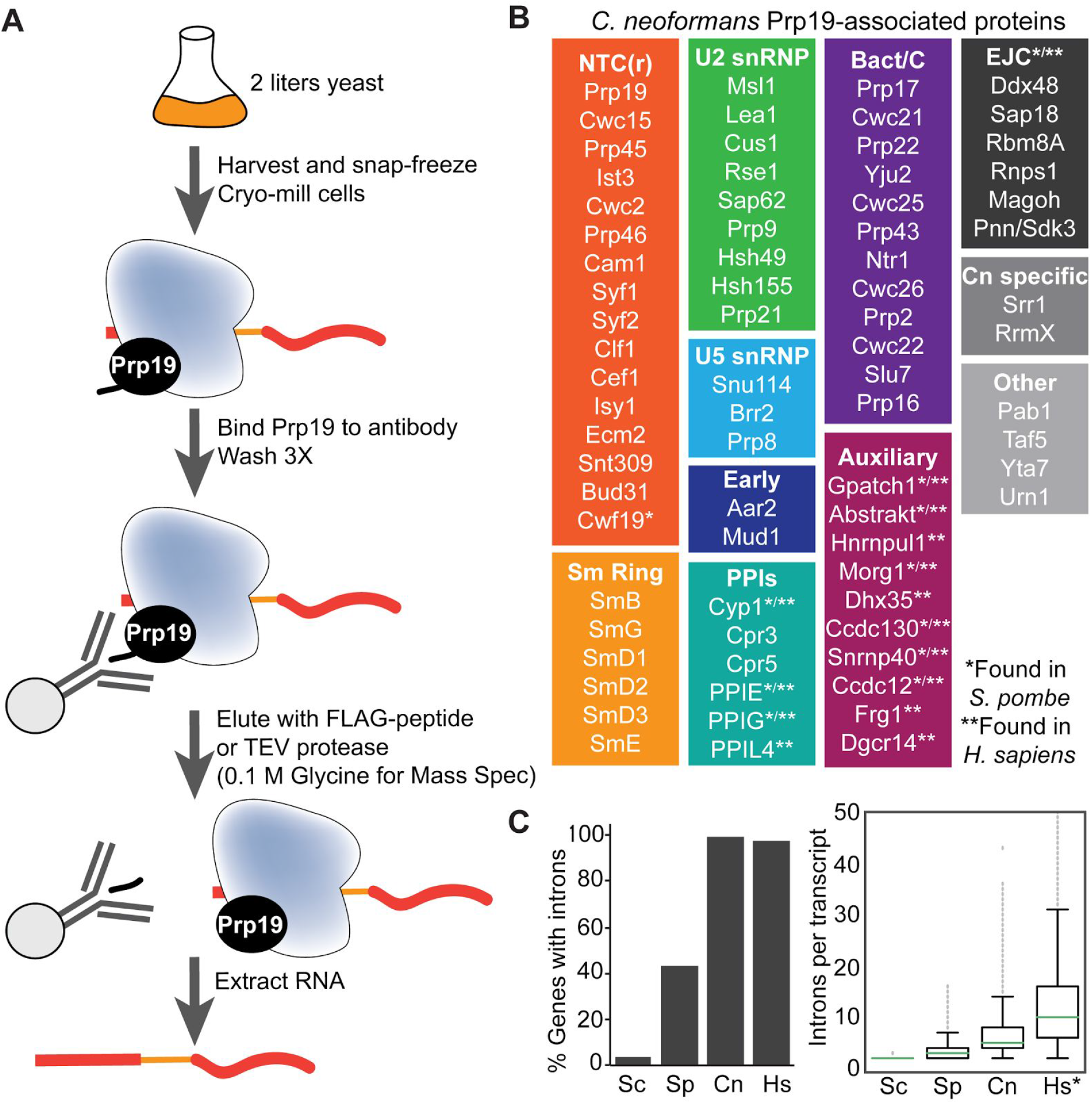
Composition of affinity purified spliceosomes, related to Figure 1. A. General scheme for purifying Prp19-associated spliceosomes (see methods for details). B. Splicing factors detected by TMT-MS in Prp19-IPs organized by complex named based on their *S. cerevisiae* orthologs, unless they are not found in *S. cerevisiae* (“Auxiliary”, “PPIs” and “EJC”) in which case they are named for the *S. pombe* or human ortholog. Category name abbreviations: NTC(r): Prp19 complex and Prp19 complex related; PPIs: peptidyl prolyl isomerases; EJC: exon junction complex; Cn specific: specific to *C. neoformans*. C. Proportion of genes with annotated introns and density of introns per transcript in each organism from this study (Sc: *S. cerevisiae*, Sp: *S. pombe*, Cn: *C. neoformans*) as well as humans (Hs).

**Figure S2.**
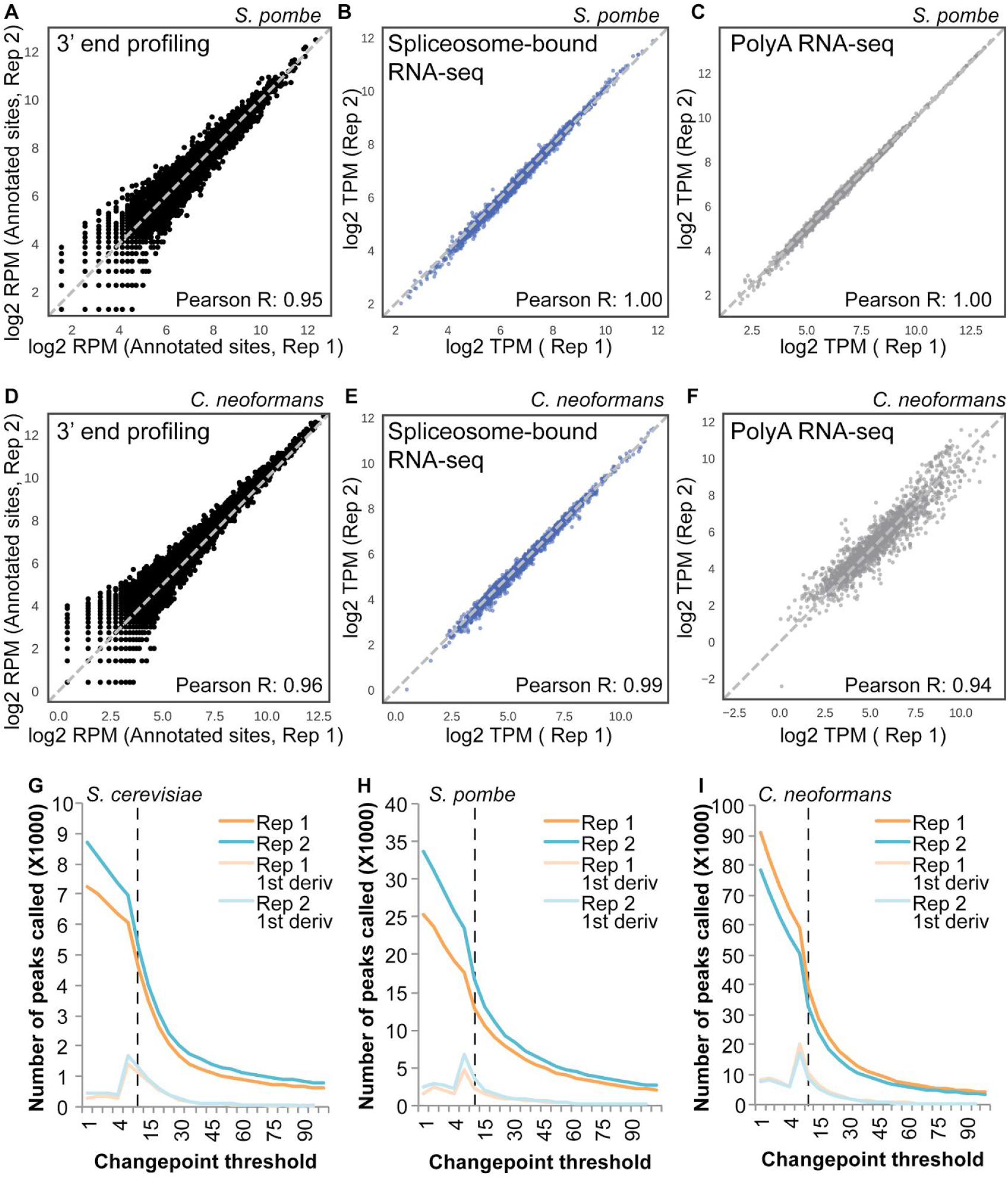
Data selection and reproducibility, related to Figure 2. Reproducibility and distributions of 3’ end profiling peak heights (A) and RNA-seq of spliceosome-bound RNA (B) and polyA RNA (C) in *S. pombe*. 3’ end profiling is measured in reads per million aligned reads. RNA-seq metrics are measured in transcript per million (TPM). D-F. Same as A-C but for *C. neoformans*. G-I. Determination of changepoint thresholds in all three yeast. Peaks were detected using changepoint analysis (see Methods) in each yeast. The number of peaks at each threshold was determined and the threshold that accomplished the largest drop in the number of peaks (maximum of the 1^st^ derivative traces, light orange and green) was chosen as the threshold.

**Figure S3.**
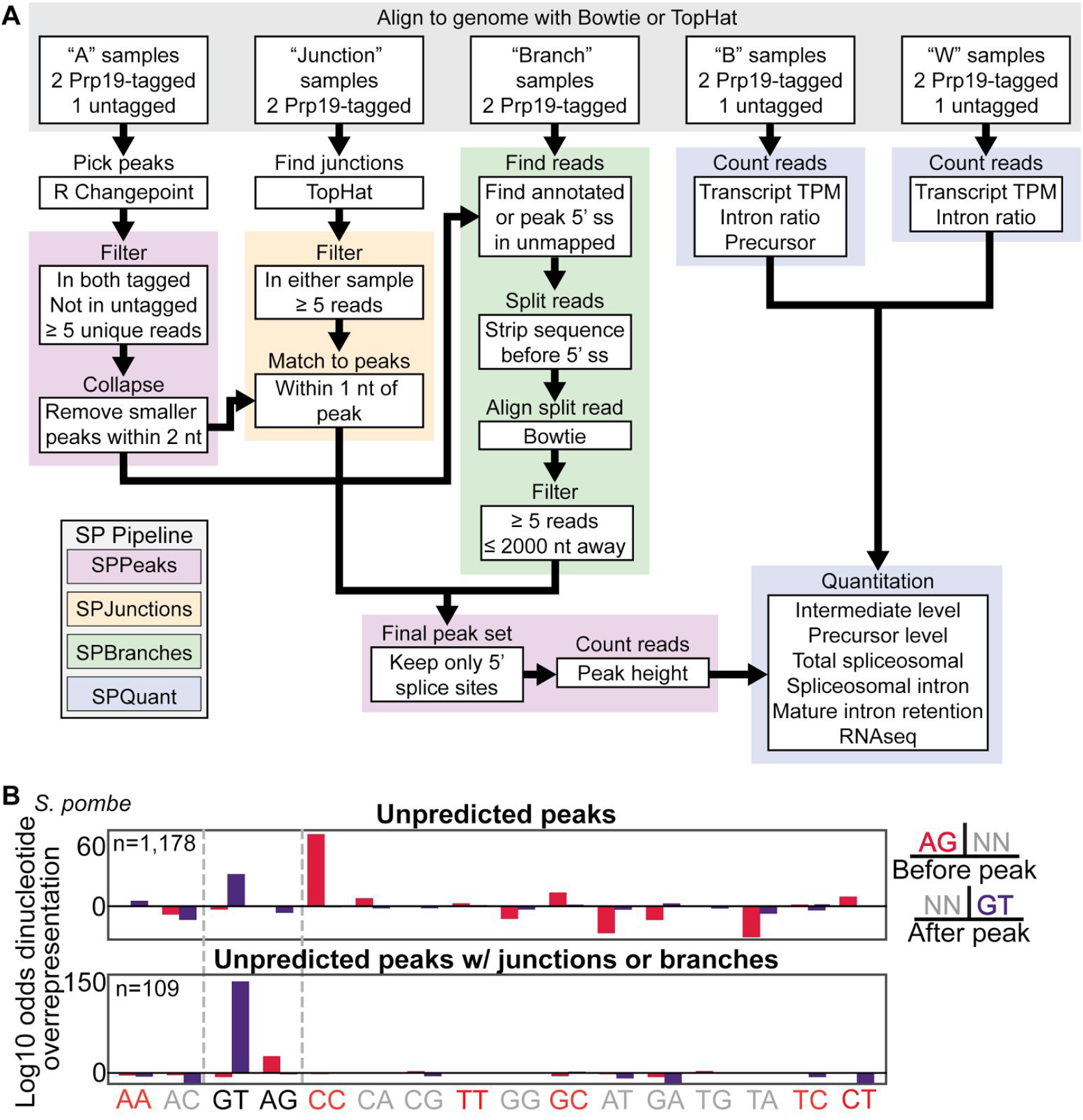
Selection of high confidence splicing events, related to Figure 3. A. Workflow and structure of the spliceosome profiling software package (SP Pipeline). See methods for additional details. B. Dinucleotide overrepresentation in unpredicted peaks and unpredicted peaks that overlap with junctions or branches in *S. pombe*.

**Figure S4.**
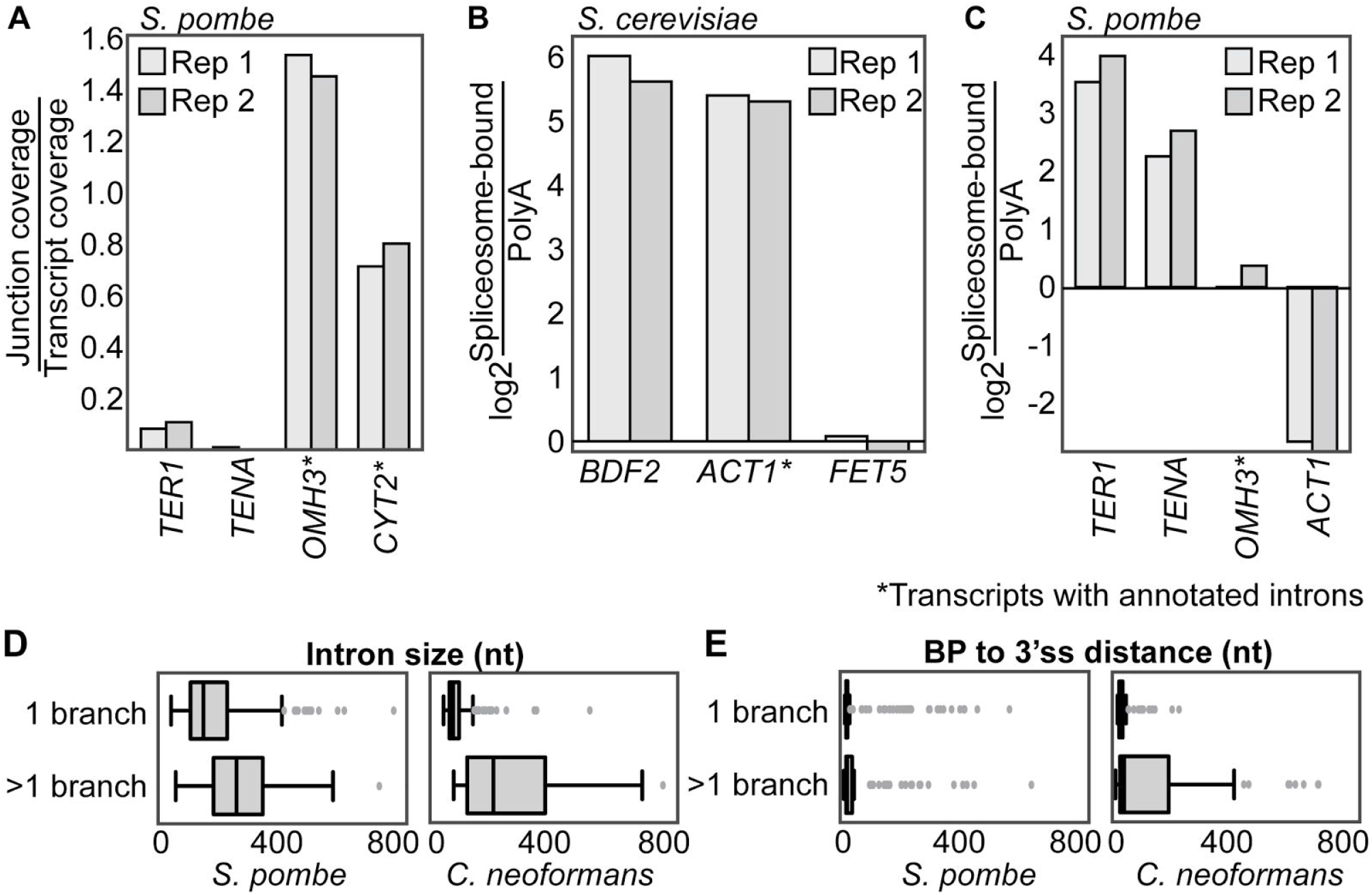
Additional information on non-canonical splicing, related to Figure 4. A. Ratio of junction coverage to transcript coverage for exon-exon junction reads originating from either interrupted splicing events (*TER1* and SPBC530.07c) or canonical splicing events (*OMH3* and *CYT2*) in *S. pombe*. B-C. Abundance of spliceosome-bound vs. polyA mRNA for transcripts with interrupted splicing events compared to canonically spliced and unspliced transcripts. D. Size distributions of introns with only one detected branch or more than one detected branch. Branches were detected with the SPBranch module of the SP Pipeline as described in the methods. E. Distance between the branch point and 3’ss in introns with only one detected branch or more than one detected branch.

**Figure S5.**
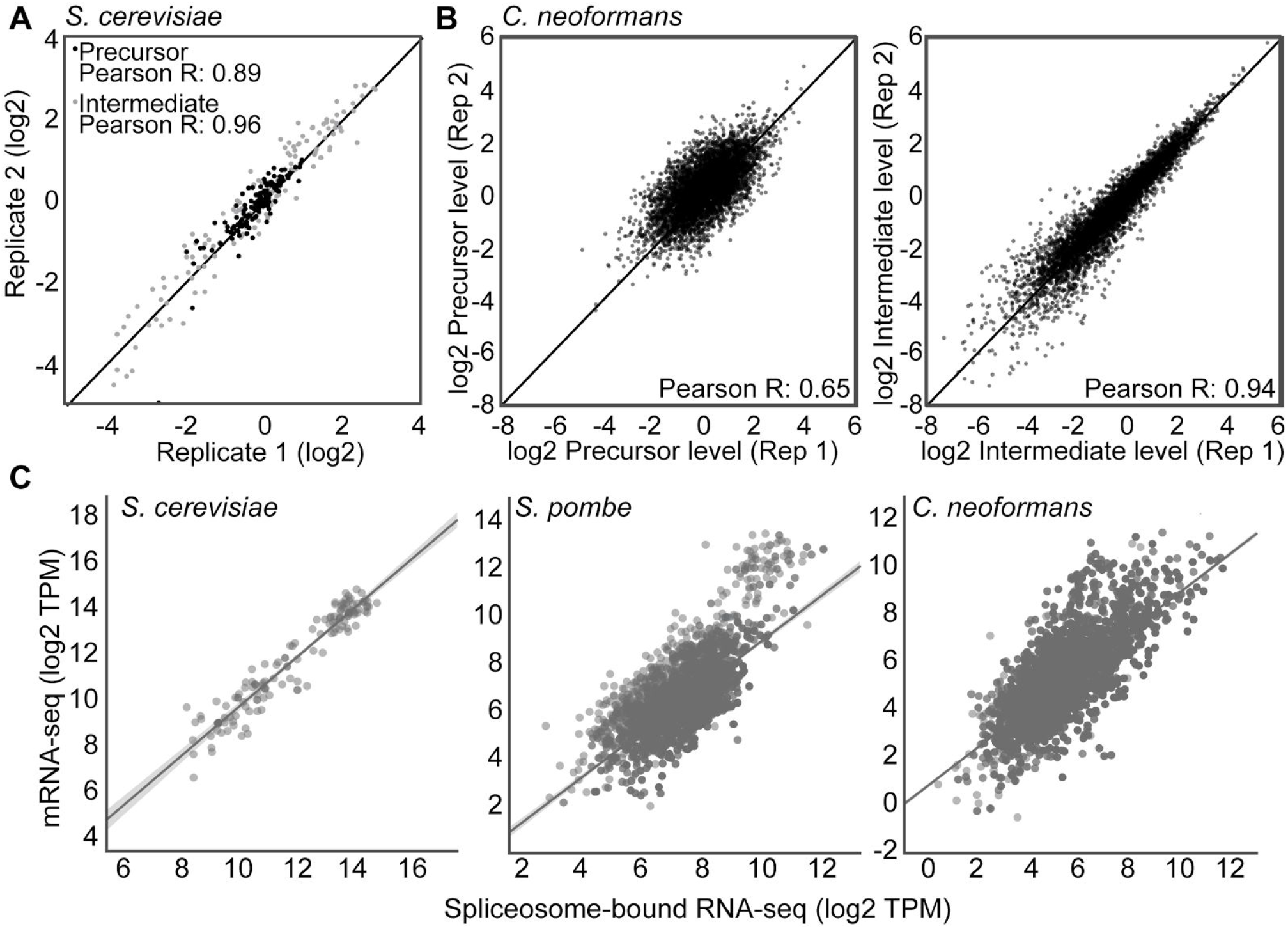
Reproducibility and distribution of precursor and intermediate levels, related to Figure 5. Distribution of precursor (dark grey) and intermediate (light grey) levels for *S. cerevisiae* (A) and *C. neoformans* (B). C. Scatter plots of the relationship between polyA RNA-seq and spliceosome-bound RNA-seq for each transcript.

**Figure S6.**
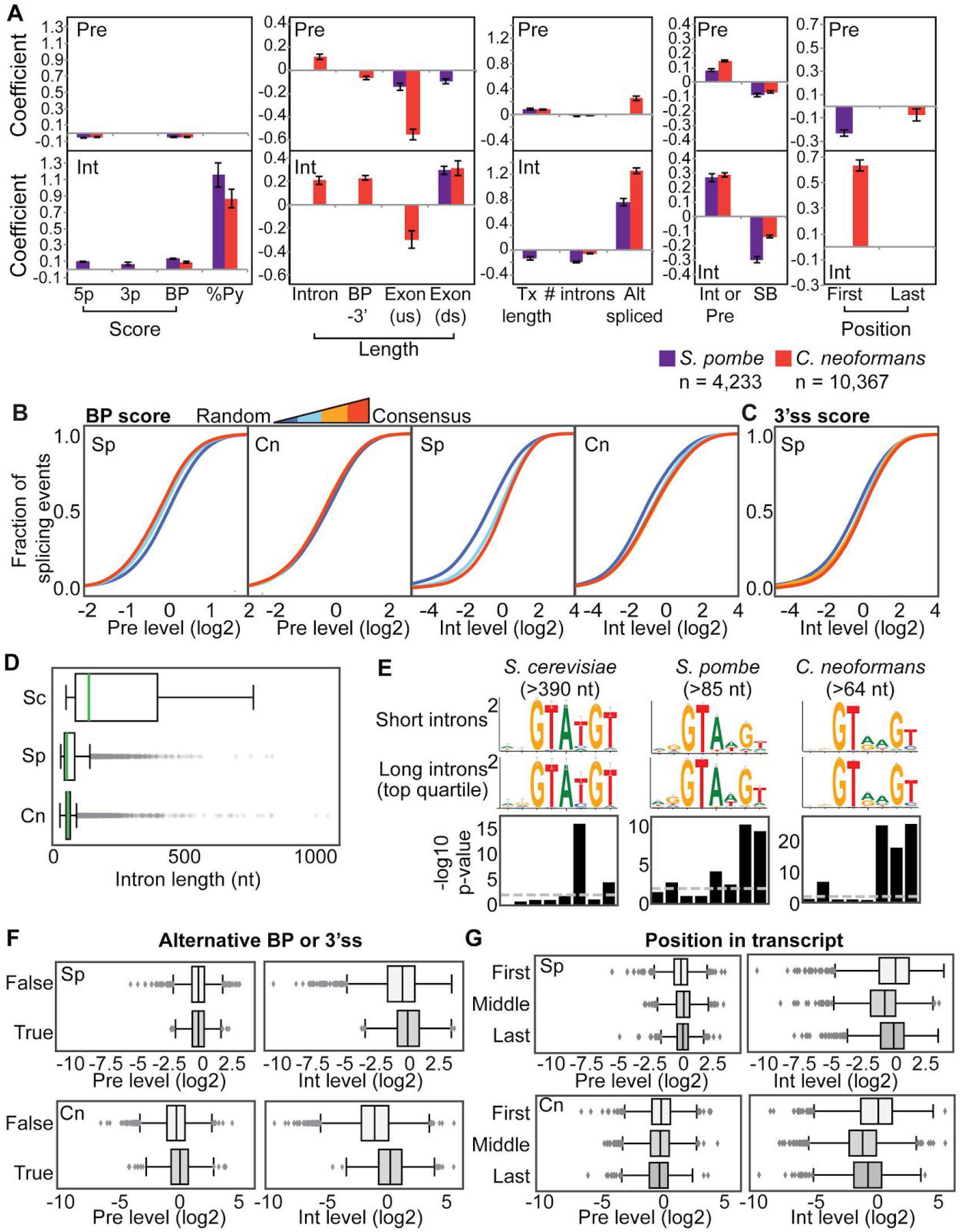
Uncovering intron features that predict splicing efficiency, related to Figure 6. A. Bayesian modeling of the correlation between intron features and precursor (Pre) or intermediate (Int) level. Coefficients for explanatory variables with a posterior inclusion probability (PIP) of at least 0.5 are shown. Error bars indicate the uncertainty of each coefficient in the final averaged model. B-C. Relationship between precursor and intermediate levels and BP or 3’ss scores. Introns were split into quartiles based on BP score (blue=random sequence, red=consensus) and the distribution of each metric is plotted as a CDF. D. Differences in intron length distributions in the three yeasts used in this study. E. Long introns have 5’ splice sites that are closer to consensus. P-values were determined by the chi-squared test for independence and logos were generated with WebLogo3 (Crooks, 2004). F-G. Difference in precursor and intermediate levels based on the whether an alternative BP or 3’ss was detected for an intron and the position of the intron in the transcript.

**Figure S7.**
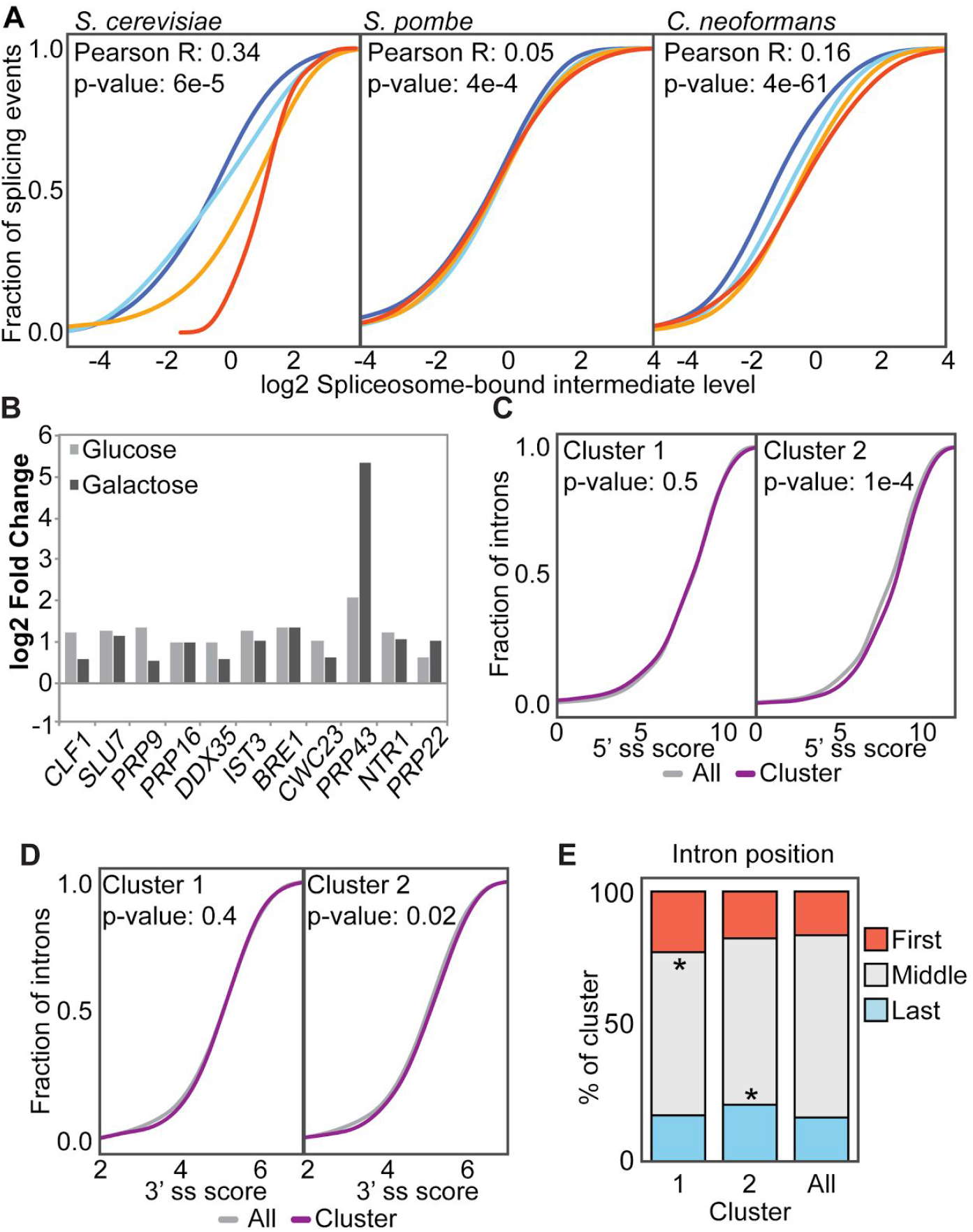
Discard of native transcripts, related to Figure 7. A. Relationship between intron retention in polyA mRNA and intermediate level. Introns were split into quartiles based on the level of intron retention calculated in polyA RNA-seq data. The distribution of intermediate level in each quartile is plotted as a CDF. B. All splicing factors that are significantly differentially expressed in *prp43DN* by polyA RNA-seq are increased (log2 fold change and significance (adjusted p-value) determined using DESeq2). C-D. 5’ and 3’ splice site scores as determined by scoring against a PSSM for each cluster (see Methods). P-values determined by Mann-Whitney U test. E. Intron position proportions of each cluster (see Figure 7). The proportion of introns that are first, last or in the middle of the transcript are plotted for the clusters with either increased precursor (cluster 1) or increased intermediate (cluster 2) vs. all introns. P-values determined by proportion hypothesis test (*p-value < 0.01).

## METHODS: KEY RESOURCES TABLE

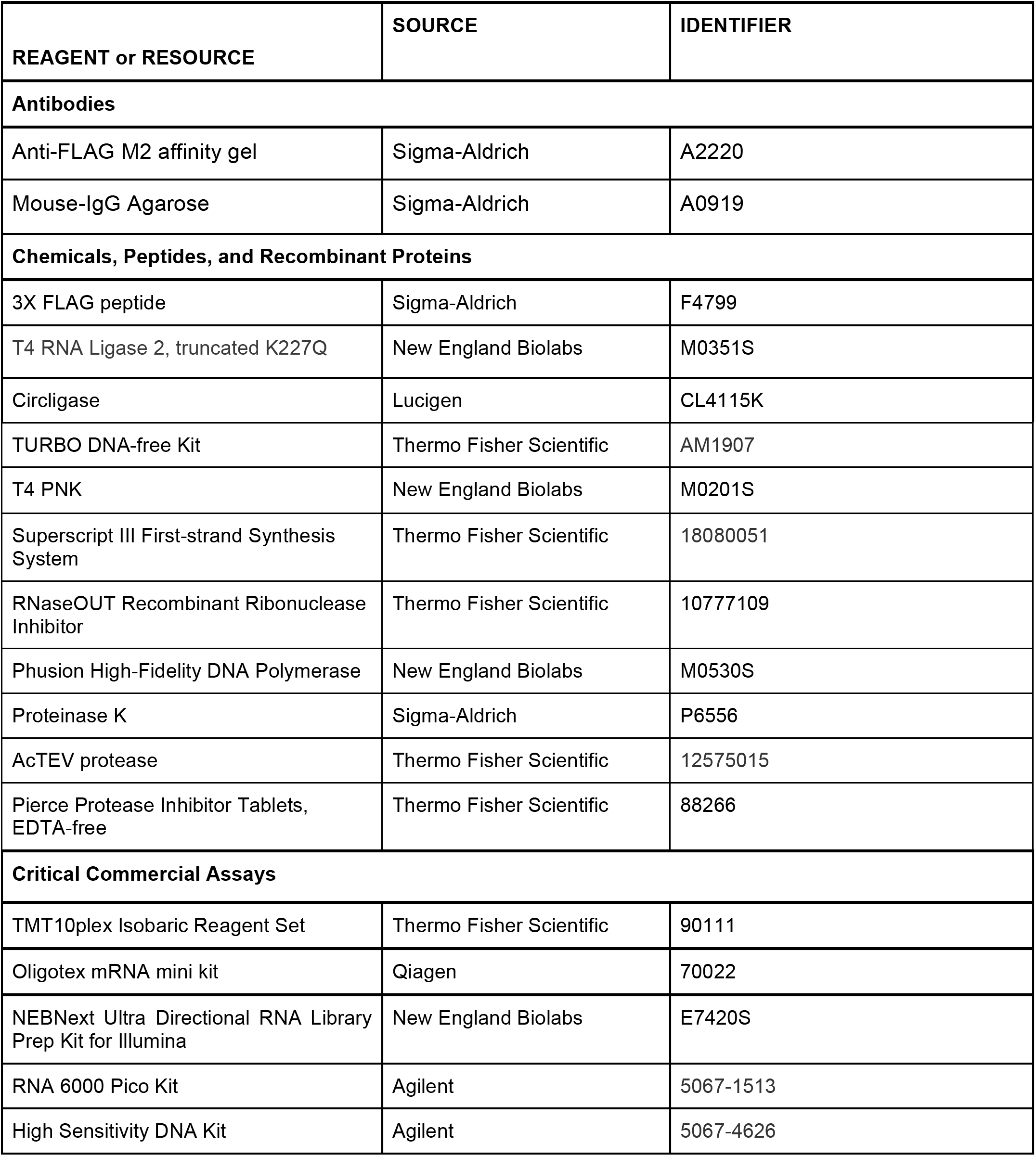
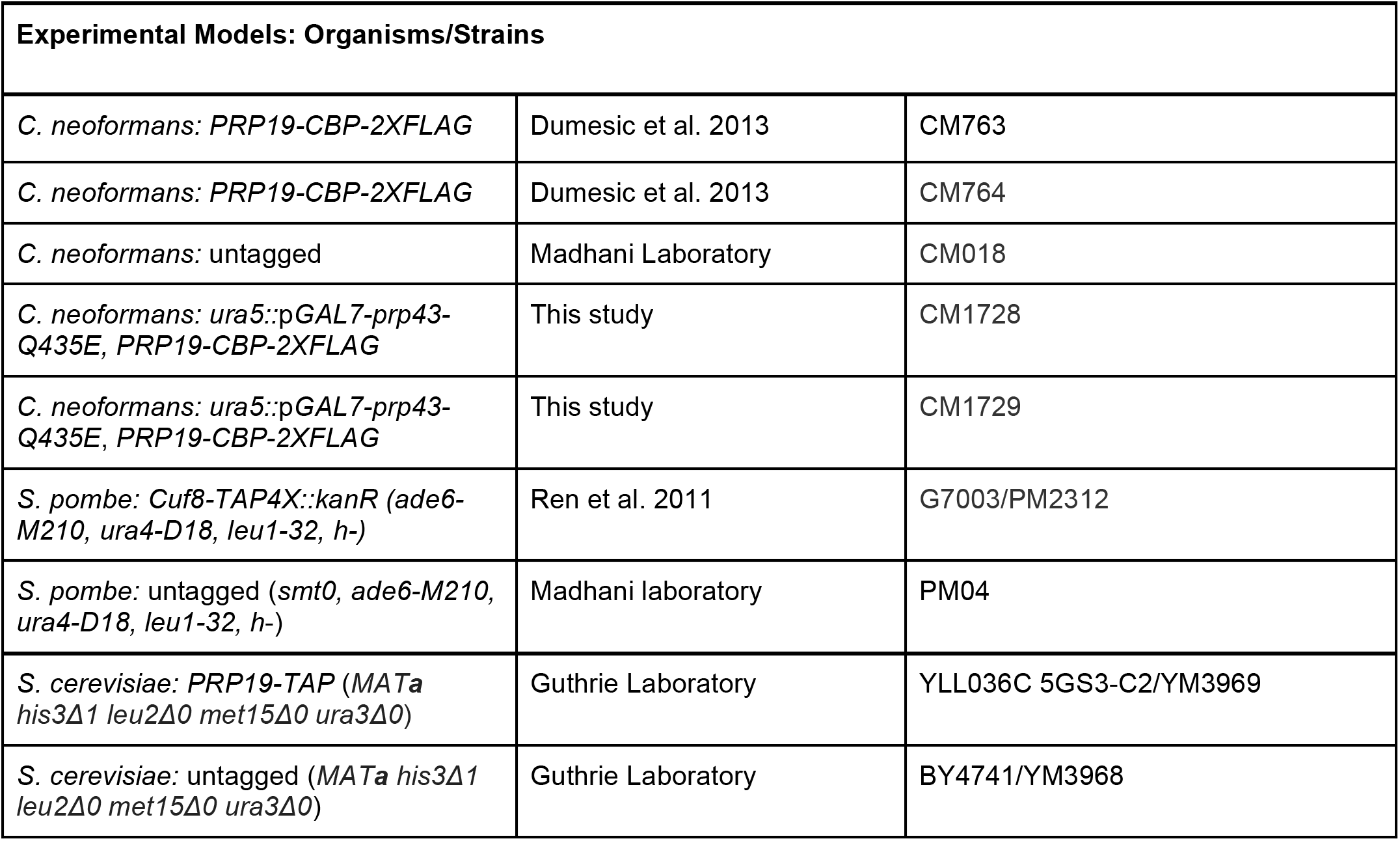
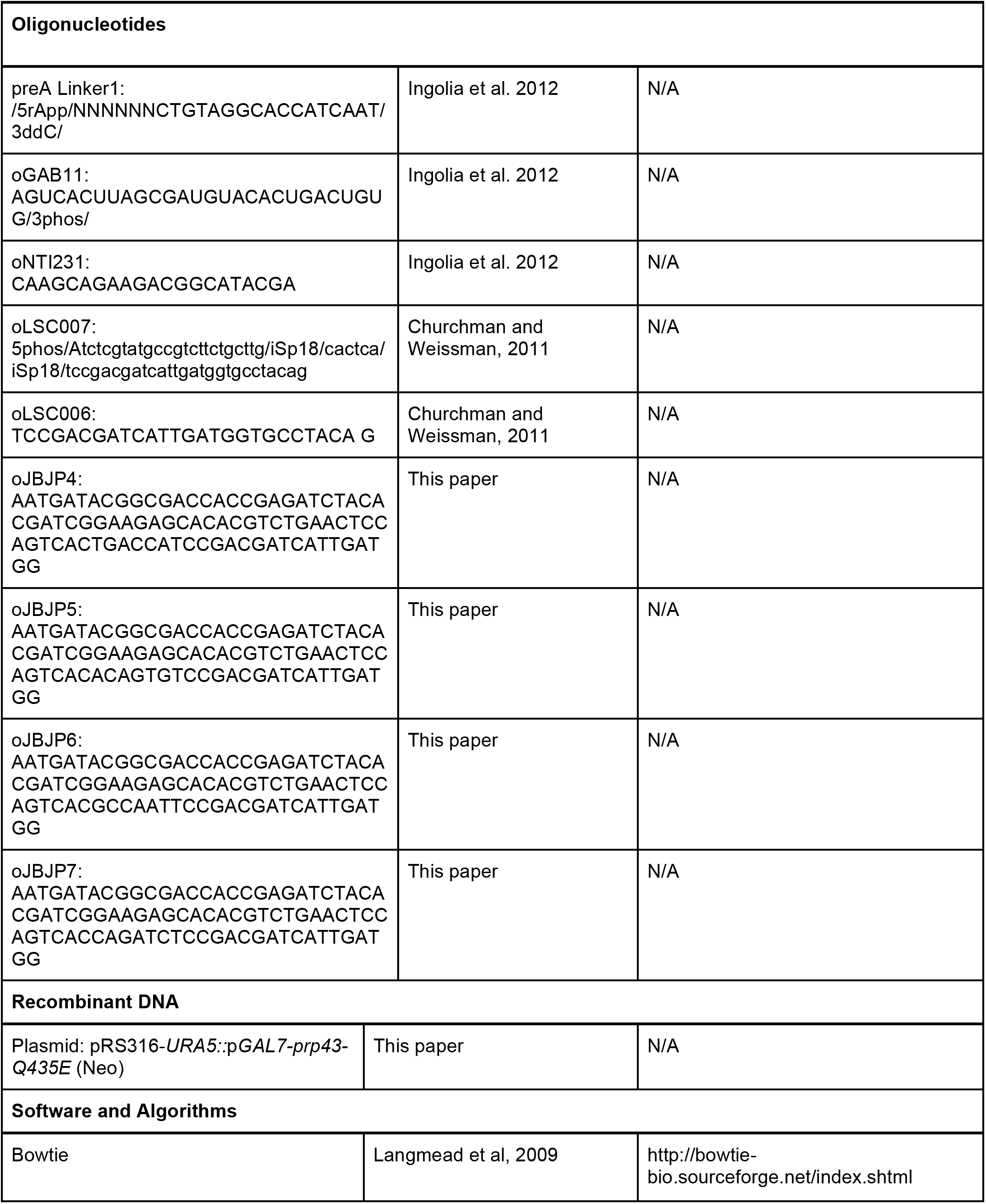
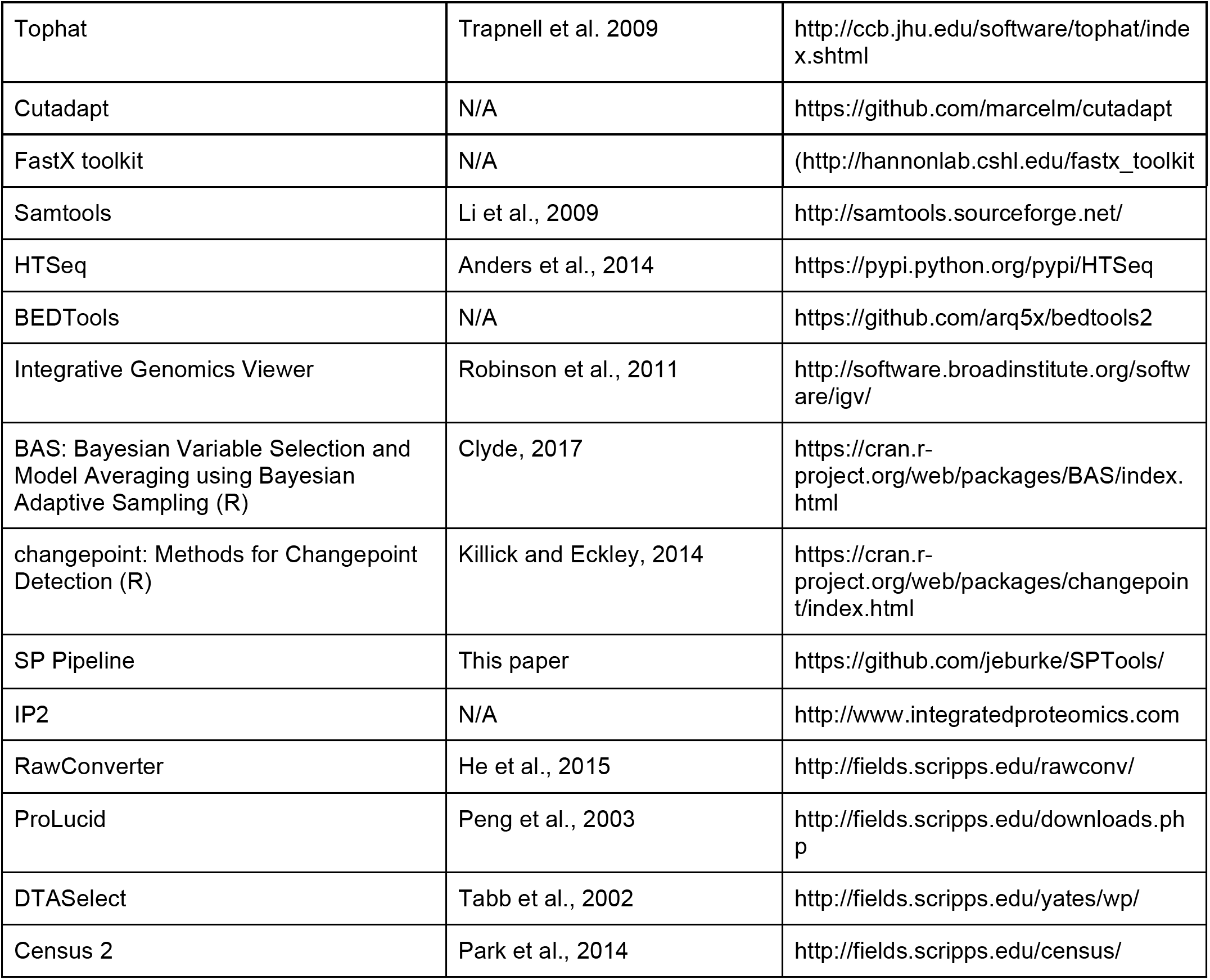

### Experimental procedures

#### Strain construction

*C. neoformans* strains were constructed by biolistic transformation (Chun and Madhani, 2010). Briefly, the ORF, 3’ UTR and downstream 500 bp of PRP43-Q435E followed by a selection marker and flanked by 1 kb of homology to the *URA5* locus were cloned into vector pRS316 using homologous recombination in *S. cerevisiae*. 10 μg of the plasmid were linearized and desiccated and then precipitated onto gold beads using spermidine and CaCl**2**. DNA-coated gold beads were shot into cells using a gene gun (BioRad, Model PDS-1000/He Biolistic-Particle Delivery System). Insertion of the full construct into the genome was confirmed by colony PCR using primers outside the sequence included in the plasmid and failure to grow on SC –Ura medium. The existence of the wild type and mutant copies of *PRP43* were confirmed by Sanger sequencing and detected by RNA-seq. Tags in all strains (including those previously constructed) were confirmed by Western blot.

#### Prp19 immunoprecipitation

2 L of all strains were grown in the appropriate rich medium (*C. neoformans* and *S. cerevisiae*: Difco YPAD, *S. pombe*: YS + 3% glucose) starting at a low density (~0.002 OD/ml) overnight at 30°C. 1% glucose was added when the cultures reached 1 OD. Cells were harvested at 2 OD by centrifugation at 5000 rpm for 6 min at 4°C. They were then washed in ice-cold sterile water and harvested again at 6000 rpm for 6 min at 4°C. Cells were resuspended in 25 ml cold H03 buffer (25 mM HEPES-KOH, pH 7.9, 100 μM EDTA, 500 μM EGTA, 2 mM MgCl**2**, 300 mM KCl (200 mM for *S. cerevisiae and S. pombe)*, 16% glycerol, 0.1% Tween) with protease inhibitors (Complete-EDTA free, Pierce) and 1 mM DTT. The resuspension was dripped into liquid nitrogen to form popcorn. Cells were lysed by cryo-grinding in a SPEX Sample Prep 6870 for 3 cycles of 2 min at 10 cps with 2 min of cooling between each cycle.

Lysate was thawed quickly with mixing in a 30°C water bath until just liquid. 500 units of RNaseOUT (Invitrogen) were added immediately after thawing. Lysate was cleared by centrifugation at 17000 rpm for 40 min at 4°C (Beckman JA17.0 rotor). Whole cell extract samples were removed and flash frozen at this point. 400 μl agarose beads (M2-FLAG (Sigma) or IgG (Sigma)) were washed in 10 ml H03 buffer three times. Lysate was incubated with the prepared beads for 3 hours at 4°C. The beads were pelleted by centrifugation at 1200 rpm for 2 min and then equilibrated by resuspending three times for 10 min with 15 ml H03 buffer (protease inhibitor in the first wash). In strains with FLAG, Prp19 was eluted three times in 150 μl elution buffer (25 mM HEPES-KOH, pH 7.9, 2 mM MgCl_2_, 300 mM KCl, 20% glycerol) with 1 mg/ml 3X FLAG peptide (Sigma). In strains with TAP, Prp19 was eluted in 500 μl 1X TEV buffer with 50 units AcTEV protease (Invitrogen) and 40 units RNaseOUT.

#### Mass Spectrometry

Immunoprecipitation of Prp19 was performed as described above, except that protein was eluted three times in 0.1 M glycine HCl, pH 3.5, for 5 minutes at 4°C. The samples were dissolved in 60 μL of 8 M urea in 100 mM TEAB, pH 8.0. Reduction was performed with at 5 mM TCEP 20 min at room temperature. Alkylation followed at 55 mM 2-chloroacetamide for 20 min in the dark at room temperature. The samples were precipitated in cold acetone (1:6) at -20 °C overnight. Tryptic digestion and Ten-plex TMT labeling (Thermo Scientific) were performed according to the manufacturer’s instructions. The ten samples were combined and run. Additionally, the five control (untagged) samples were combined and analyzed as were five experimental (Prp19-2XFLAG) conditions. The TMT labeled samples were analyzed on a Fusion Orbitrap tribrid mass spectrometer (Thermo Fisher Scientific) with multinotch data acquisition (McAlister et al., 2014). Samples were injected directly onto a 30 cm, 75 μm ID column packed with BEH 1.7 μm C18 resin (Waters). Samples were separated at a flow rate of 400 nL/min on a Thermo Easy-nLC 1000 (Thermo Fisher Scientific). Buffer A and B were 0.1% formic acid in water and acetonitrile, respectively. A gradient of 0-10% B over 20 min, 10-45% B over 270 min, an increase to 90% B over another 60 min and held at 90% B for a final 10 min of washing was used for 360 min total run time. Column was re-equilibrated with 20 μL of buffer A prior to the injection of sample. Peptides were eluted directly from the tip of the column and nanosprayed directly into the mass spectrometer by application of 2.5 kV voltage at the back of the column. The Orbitrap Fusion was operated in a data dependent mode. Full MS1 scans were collected in the Orbitrap at 120k resolution. The cycle time was set to 3 s, and within this 3 s the most abundant ions per scan were selected for CID MS/MS in the ion trap. MS3 analysis with multinotch isolation was utilized for detection of TMT reporter ions at 60k resolution. Monoisotopic precursor selection was enabled and dynamic exclusion was used with exclusion duration of 30 s.

Protein and peptide identification and protein quantitation were done with Integrated Proteomics Pipeline - IP2 (Integrated Proteomics Applications, Inc., San Diego, CA. http://www.integratedproteomics.com/). Tandem mass spectra were extracted from raw files using RawConverter (He et al., 2015) and were searched against a *C. neoformans* protein database (http://www.broadinstitute.org) with reversed sequences using ProLuCID (Peng et al., 2003; Xu et al., 2015). The search space included all half- and fully-tryptic peptide candidates, with static modifications of 57.02146 on cysteine and of 229.1629 on lysine and the N-terminus. Peptide candidates were filtered using DTASelect, with these parameters -p 1 -y 2 --trypstat --pfp .01 --extra --pI -DM 10 --DB --dm -in -t 1 --brief --quiet (Tabb et al., 2002). Quantitation was performed using Census (Park et al., 2014).

#### RNA preparation

RNA was extracted from affinity purified spliceosomes and whole cell extract by first treating with 120 μg Proteinase K in reverse buffer (10 mM Tris-HCl, pH 7.4, 5 mM EDTA, 1% SDS). RNA was then isolated by phase separation with 1 volume acid Phenol:Chloroform (1:1) and then washed with chloroform. RNA was precipitated with 2 μl GlycoBlue in 0.3 M sodium acetate and 50% isopropanol, then recovered by centrifugation for 30 min at 14000 rpm at 4°C and washed with 70% ethanol. RNA was initially checked by RT-qPCR and later by Bioanalyzer for enrichment of U2, U5 and U6 snRNAs after DNase treatment with 6 units TURBO DNase I (Invitrogen) and purification using the Zymo RNA Clean and Concentrate kit. RNA from whole cell extract was polyA selected with OligoTex beads (Qiagen) and DNase treated in the same manner as the spliceosome-bound RNA.

### 3’ end profiling and total spliceosomal RNA-seq

#### Library preparation

RNA from each IP was split with 4 parts used for the “A” sample (3’ end profiling) and 1 part for the “B” sample (spliceosome-bound RNA). “A” sample RNA was ligated to a pre-adenylated adaptor with a 5’ random hexamer (see Key Resources Table) using T4 RNA ligase 2, truncated, K227Q (NEB) for 2.5 hours at 37°C (25% PEG-8000, 12.5% DMSO, 1X buffer). After precipitation and resuspension, RNA was hydrolyzed in hydrolysis buffer (100 mM sodium carbonate, pH 9.2) for a predetermined amount of time (10-15 min) at 95°C then immediately neutralized with 0.3 M sodium acetate and precipitated. “B” sample RNA was first hydrolyzed (see “A” sample) and precipitated. Hydrolyzed RNA ends were then repaired by treating with 25 units PNK (NEB) for 1 hour at 37°C. After precipitation, “B” sample RNA was ligated to the pre-adenylated adaptor as for the “A” sample. PolyA selected samples were treated the same as the “B” samples. All samples were loaded on a 10% polyacrylamide gel with 8 M urea and 1X TBE in 1X formamide dye with bromophenol blue. The gel was run for 50 min at 200 V then stained for 5 min in 1X Sybr Gold (Invitrogen), 1X TBE. RNA bands were excised between 50 and 400 nt. RNA was extracted from crushed gel in nuclease free water for 10 min at 70°C and then separated from the gel by centrifugation in a Spin-X column (Costar) for 3 min at 13000 rpm. RNA was precipitated with GlycoBlue and 0.3 M sodium acetate in 50% isopropanol.

All samples were converted to cDNA using the Superscript III kit (Invitrogen) and the oLSC007 reverse transcription primer (see Key Resources Table). cDNA was separated from primer by denaturing PAGE (same conditions as for RNA above). Resuspended cDNA was circularized with the Circligase kit (Epicentre). Circularized cDNA was amplified by PCR with the Phusion polymerase kit (NEB) using one indexed primer and oNT231 (see Key Resources Table). Libraries were size selected on 8% non-denaturing polyacrylamide gels and confirmed by Bioanalyzer (High Sensitivity DNA kit). Libraries were sequenced on a HiSeq 4000 with single-end 50 bp reads with the oLSC006 custom sequencing primer.

#### Data analysis

5’ adaptor sequence was trimmed from the 3’ end of reads using custom scripts. Reads shorter than 24 bp after adaptor trimming were excluded from analysis. Reads were then collapsed using fastx_collapser (http://hannonlab.cshl.edu/fastx_toolkit/). The 6 nt barcode was removed from the beginning of collapsed reads using a custom script. Collapsed, trimmed reads were first aligned to rRNA and snRNA sequences using Bowtie (Langmead et al., 2009) with the following parameters:

> *bowtie -p10 -v2 -m1 --un CM764no_A_rRNA_un /home/jordan/GENOMES/crypto_rRNA -f CM764no_A_S64_L007_R1_001_debarcoded.fasta --sam CM764no_A_rRNA_al.sam*

Unaligned reads were then aligned to the genome with Bowtie using the following parameters:

> *bowtie -p10 -v2 -M1 --best --max CM763db_A_multi --un CM763db_A_un /home/jordan/GENOMES/Crypto_for_gobs -f CM763db_A_UBC4_un --sam CM763db_A_al. sam*

Conversion to BAM format, sorting and indexing was accomplished using Samtools (Li et al., 2009). BAM files were converted to bedgraph files using BEDTools (https://github.com/arq5x/bedtools2). Data were visualized using IGV (Robinson et al., 2011). Total reads in transcripts in RNA-seq style libraries were determined using custom scripts written around HTseq (Anders et al., 2014). Reads spanning 5’ splice sites (precursor) and reads beginning at 5’ splice sites (intermediate) were counted using custom scripts using the Pysam package (https://github.com/pysam-developers/pysam). Precursor values are included for sites with at least 20 reads covering the exon-intron junction.

Peaks were identified using the PELT (Pruned Exact Linear Time) algorithm (Killick et al., 2012) in the R package changepoint (Killick and Eckley, 2014). First, 5’ ends of aligned reads were converted to the bedgraph format using BedTools. Then, a range of thresholds between 1 and 100 were tested for each sample from each yeast. Peaks were detected after the threshold at which the number of detected peaks dropped most steeply (10 for all samples Figure S2G-I) was applied. Peaks called in data from both tagged samples but absent from untagged data were selected for further analysis.

### Junction and branch profiling

#### Library preparation

RNA recovered by affinity purification of Prp19 was treated with 10 units RNase R (Epicentre) or mock treated for 1 hour at 37°C. RNA was then immediately purified using the Zymo RNA Clean and Concentrate kit and DNase treated as above. Libraries were prepared using the NEBNext directional kit (NEB) according to the manufacturer’s instructions. To improve read-through of branches in lariat RNA species, we added 1 mM MnCl_2_ to the reverse transcription reaction (Madhura Raghavan and Jeffrey Pleiss, personal communication). Final size selection was performed using an 8% nondenaturing 1X TBE polyacrylamide gel. Libraries were sequenced on a HiSeq 4000 with paired-end 100 bp reads.

#### Data analysis

Adaptor was trimmed from either end of reads using Cutadapt (https://github.com/marcelm/cutadapt). Reads shorter than 25 bp after trimming or with a quality score less than 10 were not considered in the analysis.

> *cutadapt -a AGATCGGAAGA -A AGATCGGAAGA --trim-n -m 25 -q 10 -o CM763-RR_1_trim.fq -p CM763-RR_2_trim. fq ../FASTQ_FILES/CM763-RR_S31_L007_R1_001. fastq.gz ../FASTQ_FILES/CM763-RR_S31_L007_R2_001.fastq.gz*

Trimmed reads were aligned to the genome with Tophat

> *tophat -p 1 -o CM763-RR --library-type rf-firststrand -i 30 -1 400 -G CNA3_FINAL_CALLGENES_ 1_gobs.gff3 H99-2 CM763-RR_1_trim.fq CM763-RR_2_trim.fq*

Data were converted for visualization in IGV as described above. Junction position and read depth were extracted from junction.bed files from Tophat using custom scripts. Only junctions with at least 5 reads were included in the analysis.

Reads that did not align to the genome were searched for branches with custom Python scripts. First, a set of potential 5’ splice sites were established based on annotation and cleavage events that occur immediately before a “GC” or “GT” in the genome. Unaligned reads were extracted from the Tophat generated BAM file using BEDTools and searched for the 15 nt sequence immediately downstream of this site. All reads containing one of these sequences were split and realigned to the genome with Bowtie using the following parameters:

> *bowtie -p2 -v1 -M1 --best/home/jordan/GENOMES/Crypto_for_gobs -f Cn_ann_split.fa --sam Cn_ann_branches.sam*

The 5’ splice site position was embedded in the read name so it could be recovered later during analysis. Finally, potential branches were filtered based on the presence of an adenosine within 3 nt of the read split and required that the branch be with 1 kb of the 5’ splice site. Only branches with at least 5 reads were considered for further analysis.

### Statistical approaches

#### Splice site scoring

Splice site scores of each site were determined using a position-specific scoring matrix (PSSM) method. A matrix is constructed based on all the annotated splice sites in the genome as follows:

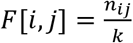 where *n_ij_* is the number of sequences with base *i* at site *j* and *k* is the total number of sequences.
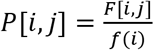 where *f*(*i*) is the frequency of the base in the genome
Finally, *S*[*i, j*] = *log*_2_(*P*[*i, j*])

Each individual splice site in a sample is scored against this matrix. Logos were generated using WebLogo3 (Crooks, 2004).

#### Branch determination

If a branch was determined experimentally, the branch with the most reads was used as the major branch site. Otherwise, branches were determined based on sequence and distance from the 3’ splice site. First, a list of all experimentally determined branches was assembled for *C. neoformans* and *S. pombe*. Each unique branch sequence was ranked based on its usage throughout the genome. Introns without experimentally determined branches were searched for each branch in this list in rank order. If no branch was found, then the adenosine closest to the 3’ splice site was chosen as the branch site. This method produced a distribution similar in terms of sequence and branch to 3’ splice site distance to experimentally determined branches.

#### Bayesian model averaging

Bayesian model averaging was performed using the R package BAS (Clyde, 2017). The optimal model was determined by Markov chain Monte Carlo sampling. Bayesian inclusion criteria (BIC) were used as the prior distribution of the regression coefficients and a uniform prior distribution was assigned over all models. Highly similar models were also obtained using traditional linear modeling.

#### K-means clustering

K-means clustering was performed using the Python package sklearn (Garreta and Moncecchi, 2013). The number of centroids was selected by the “elbow method” (Ketchen et al., 1996) by calculating the difference in the sum of squares for 10 clusters each in the range of 2 to 12 centroids.

